# EvSpark: Lossless Speculative Decoding for Hybrid DNA Foundation Models

**DOI:** 10.64898/2026.09.02.749017

**Authors:** Hao Ding, Wanchun Nie, Yuxi Zhang, Nannan Wu, Tianyi Qiu

## Abstract

Hybrid DNA foundation models combine convolutional, recurrent, and attention layers, making speculative decoding more difficult than truncating a KV cache. We present **EvSpark**, a speculative decoding system for Evo2 that verifies draft blocks in parallel and restores all three classes of inference state by selecting retained intermediate states, without replay. A compact, hidden-state-conditioned drafter proposes each block in one parallel forward pass. On Evo2 7B, a 48-prompt benchmark with three training seeds yields **2.96***×* on 43 real-sequence prompts and **3.27***×* including five synthetic controls. Acceleration persists at 262k-token context (1.84*×* –2.43*×* on two bacterial genomes) and over 32k generated tokens. Retraining the same drafter architecture for Evo2 20B and 40B yields 2.18*×* –2.46*×* on real sequences and 2.51*×* –2.78*×* on the full suite. Autoregressive drafter comparisons and batch measurements show why low draft latency, rather than acceptance alone, determines the gain. The method preserves the target distribution. Greedy checks find no non-tie divergences across 48 prompts and six checkpoints; complementary sampling diagnostics quantify decoding fidelity. A drafter trained on 30M supervised positions reaches 2.82*×* on real sequences, compared with 2.96*×* at 150M positions. In regulatory-DNA design, EvSpark achieves a median complete-workflow speedup of 1.57*×* over a calibrated batched native baseline.

## 1 Introduction

DNA foundation models span sequence analysis, design, and long-sequence generation [1, 2, 3, 4, 5, 6]. In interactive use, the latency of generating one sequence matters independently of the aggregate throughput obtainable by batching many requests. Evo2 7B illustrates this constraint: single-token decoding repeatedly reads approximately 13.2 GB of bf16 weights and reaches about 45–51 tok/s at short context in our native decode benchmark. At 262k context, attention-cache reads further increase latency.

Speculative decoding amortizes a target-model forward pass over several tokens. A lightweight drafter proposes a block, the target verifies it in parallel, and rejection sampling preserves the target distribution [14, 15]. For a Transformer, restoring the state after rejection mainly requires truncating the readable KV cache. Evo2’s StripedHyena2 architecture adds two other state classes: finite impulse response (FIR) windows and infinite impulse response (IIR) recurrences. A correct implementation must restore all three to the same accepted prefix. Replaying that prefix is possible, but adds a second target computation to each speculative round.

Previous work establishes speculative decoding for genomic Transformers (BioSpecDec [7]) and state recovery for state-space or hybrid models [8, 9, 10]. EvSpark addresses the joint FIR/IIR/KV state of StripedHyena2. Its block forward exposes the intermediate states needed to restore any accepted prefix by index selection. A DSpark-style [11] distilled drafter then exploits the nearly flat cost of verifying short blocks. The central design requirement is to make both rollback and drafting cheap enough that accepted tokens translate into wall-clock savings.

We make three contributions:

- **Replay-free verification for a hybrid target**. An initial-state block forward and a common state-slicing protocol restore FIR, IIR, and KV state consistently. An identity-drafter control improves from 0.54*×* to 1.05–1.16*×* native throughput when replay is removed.
- **A practical distilled drafter**. A parallel trunk conditions on a short target-feature window. We evaluate draft length, training budget, and feature selection, and compare it with truncated-target and independent small-model drafters. These comparisons separate acceptance quality from the latency required to obtain it.
- **Evaluation across scale and operating conditions**. The 7B benchmark reaches 2.96*×* on real sequences, with tests extending to 262k context and 32k generated tokens. Experiments on 20B and 40B establish transfer of the implementation and training recipe. Batch and multiple-candidate measurements characterize its operating limits, while token- and sequence-level checks quantify decoding fidelity. A regulatory-DNA design evaluation measures complete workflow time, predictive quality and diversity against an optimized batched baseline.

## 2 Background

### 2.1 Hybrid inference state

Evo2 7B [1] is a 32-layer StripedHyena2 model with five attention layers and 27 Hyena layers: nine each of the short-, medium-, and long-filter variants (HCS, HCM, HCL). It uses a byte-level tokenizer padded to 512 entries. Our main experiments use the 1M-context checkpoint and temperature 1.0 with top-*k*=4 sampling. The transform acts on the full vocabulary; it is not a hard-coded restriction to A, C, G, and T.

Hyena states have a fixed size for each sequence, whereas attention’s KV cache grows with context (Table 1). In particular, an HCL state obeys a modal recurrence of the form *s*_*t*_ = *λ ⊙ s*_*t−*1_ + *x*_*t*_. Its dependence on the full history prevents rollback by simply discarding recent inputs.

**Table 1.** Inference state and rollback in Evo2 7B. Hyena state is fixed in size (approximately 15 MB per sequence); KV storage grows with context.

| State | Layers | Retained information | Rollback operation |
| --- | --- | --- | --- |
| Outer FIR | 27 | Last 2 projected inputs | Select a window |
| HCS / HCM inner FIR | 9 / 9 | Last 6 / 127 filter inputs | Select a window |
| HCL IIR | 9 | Modal recurrence state | Select a per-position state |
| Attention KV | 5 | Keys and values, 82 KB/token | Reset readable-length offset |

### 2.2 Acceptance and the latency budget

A speculative round proposes *γ* tokens and accepts a prefix of length *k ≤ γ*. It emits the accepted prefix followed by a correction token, or a bonus token if all drafts are accepted. We use *τ* = *k* + 1 for the number of *emitted tokens per round*, and 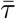 for its measured mean. A final round may be truncated by the output budget. The training proxy 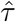 is reported separately. The approximate speedup is

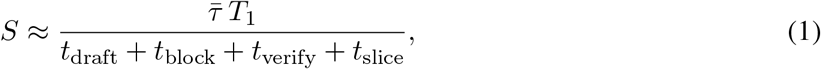

where *T*_1_ is the native per-token latency. This decomposition explains why higher acceptance need not produce higher speedup.

On the 4090, a parallel-forward probe costs about 26–27 ms for block lengths 4–16 and remains near 27 ms through length 32. Its normalized cost *T*_v_(*γ*)*/*(*γT*_1_) falls from 0.167 at *γ*=8 to 0.084 at *γ*=16. We distinguish this fixed-length probe, denoted *T*_v_, from *t*_block_ measured inside the stateful speculative loop, which verifies an anchor plus *γ* drafts. Both show the opportunity to amortize weight reads, but their timings are not interchangeable.

## 3 Method

EvSpark combines a feature-conditioned parallel drafter with block verification and state slicing in the frozen target. Figure 1 connects the network architecture to the operations performed in one speculative round.

**Figure 1:**
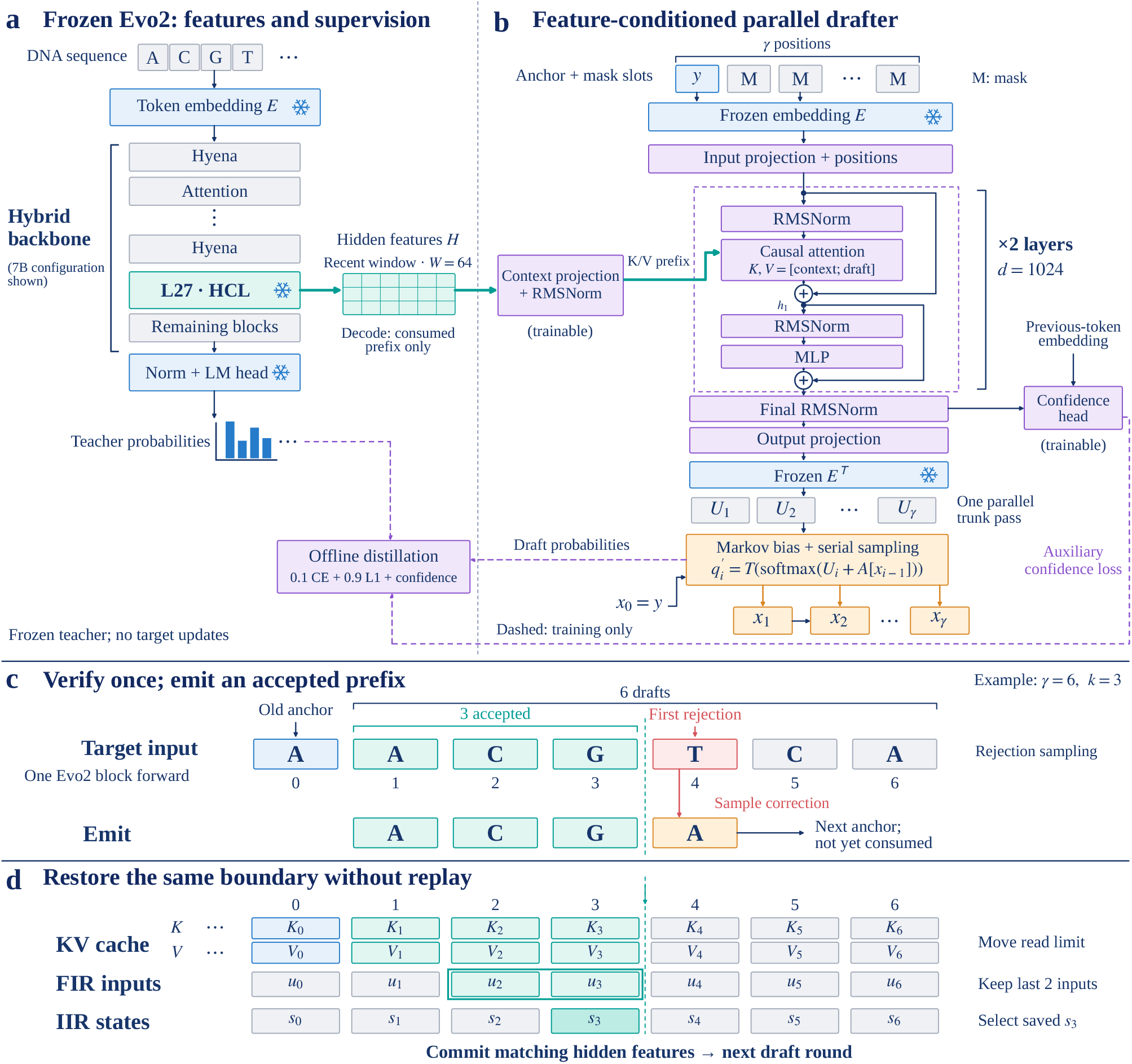
EvSpark architecture and speculative decoding. **(a)** Frozen Evo2 supplies intermediate features and offline teacher supervision; the 7B configuration extracts layer 27 (HCL). **(b)** A projected window of up to 64 target states forms the K/V prefix of each of two causal drafter layers (*d* = 1024). Anchor and mask embeddings produce all draft logits in one trunk pass; a previous-token Markov bias then supports serial sampling. Input and output embeddings are tied and frozen; projections and the two-layer trunk are trained. Dashed paths denote offline supervision using teacher-forced draft probabilities before the sampling transform *T* ; serial sampling is shown for decoding. The confidence head does not control decoding. **(c)** The same Evo2 verifies the old anchor and six drafts in one block. Three drafts are accepted; a sampled correction becomes the next unconsumed anchor. **(d)** State returns to block index 3: shorten the readable KV prefix, select the last two outer-FIR inputs *u*_2_, *u*_3_, and retain IIR state *s*_3_. The feature buffer commits the same consumed prefix for the next round; no target computation is replayed.

### 3.1 Block verification from a persistent state

At the start of a round, the target has consumed a prefix of length *L*_0_. The most recently emitted token, *y*, remains unconsumed and serves as the *anchor*. Given drafts *x*_1_, …, *x*_*γ*_, we run one block forward on [*y, x*_1_, …, *x*_*γ*_]. Logit row *i*, indexed from zero, predicts the next token after consuming *y, x*_1_, …, *x*_*i*_. This alignment supplies the target distribution for every draft and for the full-accept bonus token.

The block forward must propagate the existing Hyena state. FIR filters convolve the cached window concatenated with the new block. HCL filters combine the zero-state parallel convolution with the decaying contribution of the initial IIR state. For the modal recurrence *s*_*j*_ = *λ ⊙ s*_*j−*1_ + *u*_*j*_, the state after block position *j* ∈ {0, …, *γ*} is

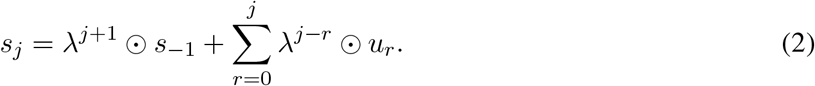

Here *s*_*−*1_ is the incoming state before consuming the anchor *y, u*_*j*_ is the modal input contribution, and powers and products involving *λ* are elementwise. The sum is the zero-initial-state response. The same computation retains the modal state at every block position in fp32 and writes the terminal state to the recurrent cache. Attention writes keys and values for the block into its existing cache.

In the evaluated Vortex implementation, the native post-prefill Hyena branch consumes only the final input token; invoking it directly on a draft block therefore violates this contract. We implement an explicit initial-state chunk branch and check its logits against sequential decoding.^1^

### 3.2 Rejection sampling and state slicing

Let 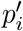 and 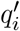 be the target and draft distributions after the chosen temperature and top-*k* transforms. At position *i*, both condition on the emitted prefix (including the unconsumed anchor) and preceding drafts *x*_1_, …, *x*_*i−*1_. Draft tokens are sampled from 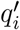 and accepted with probability 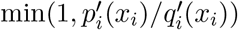. At the first rejection, the verifier draws a replacement from 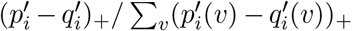 and ends the round. If all drafts are accepted, it draws a bonus from 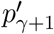 [14, 15]. The verifier must receive the distribution that actually generated each draft; using the pre-transform *q*_*i*_ in its place is incorrect. If the residual normalizer is zero, the distributions coincide and rejection has probability zero. Tokens with 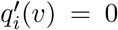 cannot be proposed. In greedy decoding, proposals are retained while they match the target argmax; the first mismatch emits the target argmax, using a fixed tie-breaking rule. Losslessness concerns the output distribution, not equality of independently sampled sequences.

After *k* accepted drafts, the target should have consumed the old anchor and those drafts, but not the newly emitted replacement or bonus. We restore the state at block index *j*^*\**^ = *k*:

- **KV:** set the readable-length offset to *L*_0_ + *j*^*\**^ + 1; entries beyond that offset remain allocated but are not read.
- **FIR:** select the last *K −* 1 filter inputs ending at *j*^*\**^ from the cached window concatenated with the block. Short histories grow according to the native recurrence and filter-tap convention until a full *K −* 1-input history is available.
- **IIR:** select *s*_all_[…, *j*^*\**^], the state already computed at that position by the block forward.

If the output budget truncates a round to *m* emitted tokens, use *j*^*\**^ = *m −* 1. If all drafts are accepted and the round is not truncated, the terminal state is already correct and slicing can be skipped. The hidden-state buffer used by the drafter commits the same consumed prefix.

This protocol removes replay. An identity-drafter control runs at 0.54*×* of native throughput with snapshot-and-replay and at 1.05–1.16*×* after slicing in two measurement campaigns. These are throughput ratios. Slicing itself takes approximately 0.62 ms; it replaces a second target forward costing about 22 ms. Figure 1c,d and Algorithm 1 summarize the resulting loop.

#### Algorithm 1

EvSpark generation with state slicing

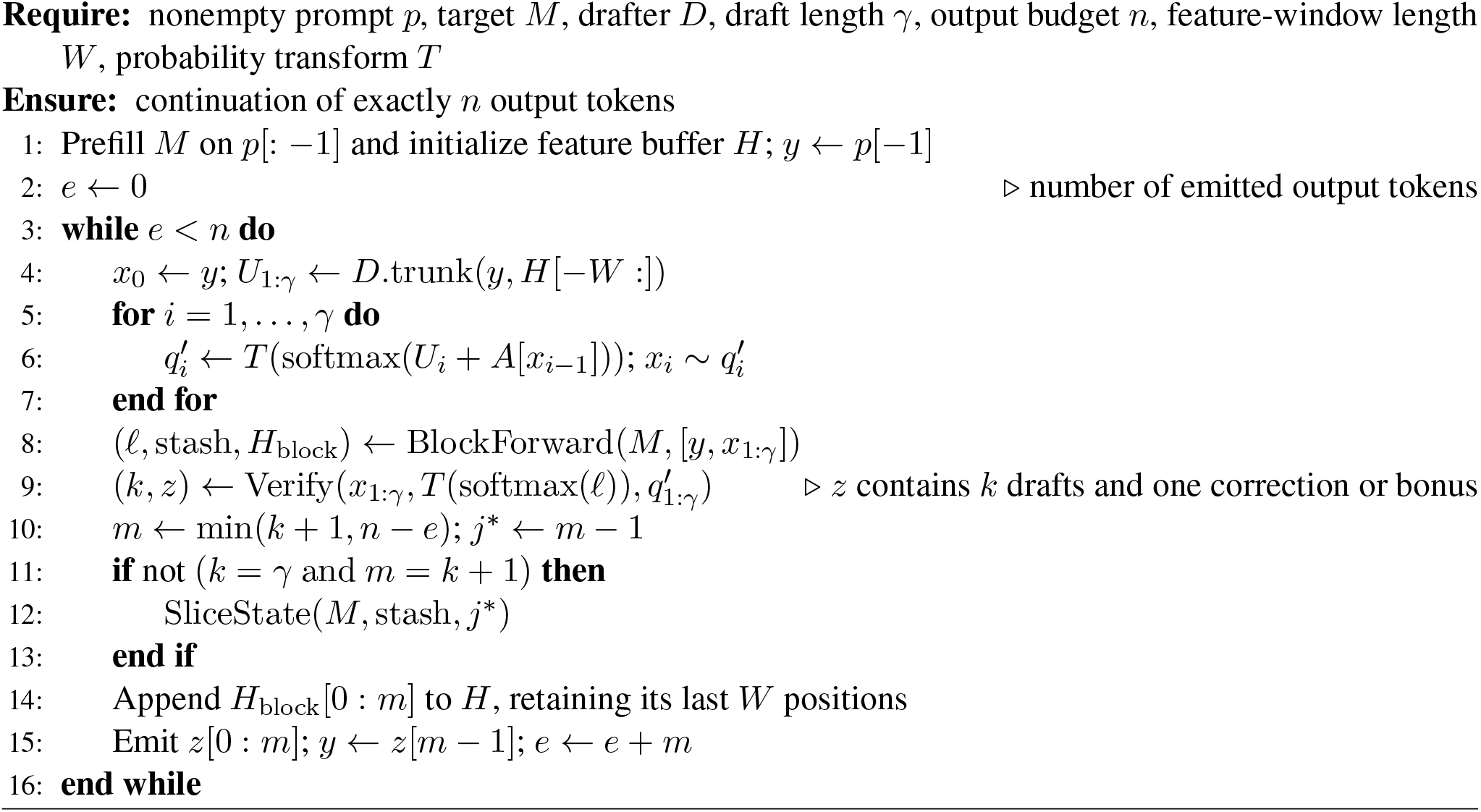

#### Algorithm conventions

*T* acts on probabilities: at positive temperature *θ*, it raises each probability to 1*/θ*, retains the top-*k* entries, and renormalizes. This corresponds to temperature-scaled logit sampling; the implementation uses numerically stable transforms. All vocabulary entries are considered before truncation. Softmax and *T* act row-wise. *A* is the learned previous-token bias. The feature slice [0 : *m*] is zero-based and half-open, whereas *x*_1:*γ*_ includes all *γ* drafts.

##### Distributional guarantee

In exact arithmetic, the block forward supplies the same conditional logits as sequential evaluation, and slicing restores the state for the same consumed prefix. Standard rejection sampling then preserves the transformed target distribution. Induction over rounds establishes sequence-level exactness (Appendix A).

##### Numerical implementation

Both native and block decoding use the deployed Evo2 7B target, which returns bf16 logits. The single-sequence verifier promotes these values to fp64 for probability transforms and rejection sampling. Block and sequential kernels can round differently near ties; Section 4.8 reports greedy, sampling, and sequence-statistic comparisons of the deployed paths.

### 3.3 A parallel, feature-conditioned drafter

We adapt the DSpark architecture [11]: a *γ*-parallel trunk with two causal attention layers of width *d*=1024 conditions on the anchor and a window of *W* =64 target hidden states (Figure 1a,b). The target features pass through a shared projection and RMSNorm, then supply a K/V prefix to each layer’s attention. Each layer uses pre-normalized attention and MLP sublayers with residual connections. The draft input consists of the anchor and *γ −*1 mask slots, with learned position embeddings. The frozen target embedding is tied between the draft input and output; trainable projections connect it to the trunk. A full-matrix Markov bias *A*[*x*_*i−*1_] conditions each position on the preceding sampled token, and a confidence head predicts acceptance. Drafting requires one trunk forward, followed by a short serial sampling loop over its outputs. The confidence head is trained but does not control scheduling in these experiments. A model trained at length *γ* supports any decode length *γ*^*′*^*≤ γ* through an exact prefix computation; extending beyond the training length is not supported. The feature-window bound applies only to the drafter: the target retains its full preceding context and recurrent state.

For 7B we capture the output of layer 27 (HCL). We select this layer based on single-layer storage cost and measured performance. At matched 30M-position budgets, several Hyena and attention injection sites give similar speedups (Section 4.7). The 20B and 40B extensions use late HCL layers at similar relative depths.

#### Offline distillation

The 7B teacher cache contains 289M recorded positions from nine sources: GTDB [30], mRNA, IMG/VR [31], eukaryotic sequence, NCBI coding sequence, ncRNA, organelles, promoters, and hg38. It stores token IDs, top-8 teacher probabilities and their residual mass, and int8 hidden states in 4096-token windows, requiring approximately 1.2 TB. Teacher targets are reconstructed by retaining and renormalizing the top four probabilities at temperature 1. Source-weighted sampling defines the intended training mixture independently of cache size (Appendix C).

For each training anchor, reference-sequence predecessor tokens condition the Markov bias at all draft positions. Let *a*_*i*_ be the observed next token in the training sequence, *p*_*d,i*_ the drafter’s untruncated softmax under this teacher-forced context, and *p*_*t,i*_ the reconstructed teacher target. The loss is

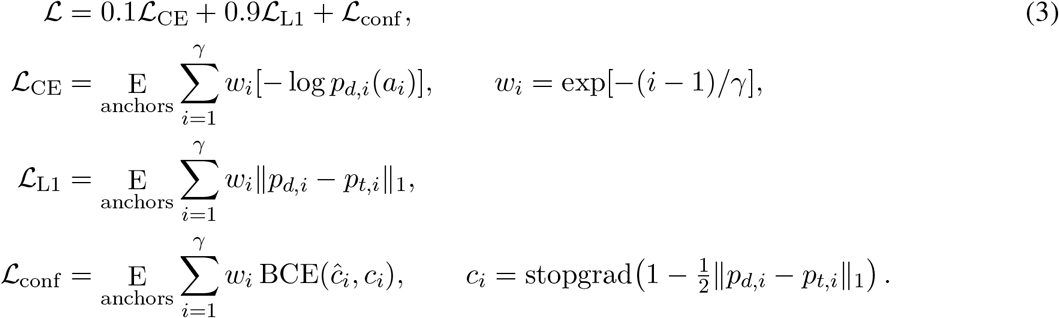

Position losses are summed with weights *w*_*i*_ and averaged over anchors. The confidence target is detached. At inference, the Markov bias conditions on the actual sampled predecessor, and the verifier receives the resulting transformed proposal distribution. Training uses AdamW with learning rate 3 *×* 10^*−*4^, cosine decay to one tenth of that value, weight decay 0.01, *β* = (0.9, 0.95), gradient clipping at 1.0, and bf16 autocast. Target parameters and tied embeddings remain frozen.

Late-layer feature collection requires care: before the final normalization, residual magnitudes in blocks 30–31 can reach 2.5 *×* 10^11^, overflowing fp16 quantization scales. We store these scales in fp32 and check for overflow. The codec belongs to the offline distillation pipeline; it does not quantize the target’s generation state.

## 4 Experiments

### 4.1 Evaluation protocol

The main benchmark comprises 48 prompts: 43 real sequences spanning bacterial coding, viral coding, human intergenic sequence, human repeats, and five OpenGenome2 domains, plus five synthetic i.i.d. controls. Each run generates 1024 tokens at temperature 1.0 and top-*k*=4, unless stated otherwise. We report the mean of per-prompt speedups, with *real-43* as the primary aggregate and *all-48* including synthetic controls. Synthetic sequences are an easier reference regime and increase the overall mean.

Speedup is native decode wall-clock divided by speculative decode wall-clock for the same prompt and output length. Timings are CUDA-synchronized and include the entire speculative loop. Both arms use the same chunked prefill, which is excluded: these are decode speedups, not full-request speedups. The 7B configuration grid uses three training seeds and a fixed decode repetition per prompt (rep-0). We average per-seed suite means; prompt-paired bootstrap analyses use 10,000 resamples. Dedicated comparisons use the replication schemes in Table 2.

**Table 2.** Replication and hardware for the main performance experiments. Every suite run has 48 prompts and 1024 generated tokens per prompt.

| Experiment | Replication | Hardware / target attention |
| --- | --- | --- |
| 7B configuration grid | 3 training seeds; rep-0 | RTX 4090 / flash attention |
| AR drafter comparison | Frozen AR weights; 1 neural checkpoint; 2 decode repetitions | RTX 4090 / flash attention for 7B |
| Multiple candidates | 1 neural checkpoint; 2 decode repetitions | RTX 4090 / flash attention |
| 20B and 40B | 1 training seed per budget; 2 decode repetitions | H20 96 GB / SDPA |
| 7B hardware retest | 1 deployment checkpoint; 2 decode repetitions | H20 96 GB / flash attention |

#### Configurations and hardware

The deployment drafter uses layer 27, *γ*=12, and 150M supervised training positions. The lower-budget alternative uses the same architecture at 30M positions. The 4090 experiments include a non-standard 48 GB card: 262k-context runs peak near 36 GB and do not fit on a standard 24 GB 4090, whereas our single-sequence experiments through 51k do. Relative to native decoding, EvSpark adds approximately 0.5 GB at 1k context and 2.1 GB at 262k, including the drafter and feature-capture buffer.

The 20B/40B experiments use their official Transformer Engine loading path and SDPA attention. A flash-attention custom-operator registration failure in the evaluated software stack prevented its use for these end-to-end runs.^2^ All H20 batch curves reported below also use SDPA; the 7B H20 end-to-end retest uses flash attention. Reported timings retain these model-specific backend choices.

#### Evaluation independence

GTDB evaluation uses accession-prefix holdouts, IMG/VR uses its validation split, and OG2-domain prompts use held-out source chunks. Prompt sequences, source records and independence annotations are supplied in Supplementary Data 1. Some hg38/chr21 and lacZ loci overlap the source records used for training, although each record has a contiguous 90%/10% train/held-out split. We therefore include 27 independent-locus control prompts in the 48-prompt suite. Region-level standard deviations describe run heterogeneity; overlapping windows should not be interpreted as independent biological replicates.

### 4.2 Nearly threefold acceleration on real sequences with Evo2 7B

The deployment configuration reaches **2.96***×* on real-43 and **3.27***×* on all-48. The respective per-seed suite means are 2.89/3.05/2.93 and 3.26/3.32/3.24. The 30M-position alternative reaches 2.82*×*/3.15*×* (real/all). Figure 2 shows that gains vary substantially by sequence class: mean speedup ranges from 2.02*×* for bacterial coding to 3.81*×* for human intergenic sequence, with synthetic controls at 5.95*×*. Independent controls give 2.33*×* on seven held-out bacterial coding prompts and 2.64*×* on two human subtelomeric windows, consistent with gains extending beyond the overlapping loci. Complete region statistics appear in Appendix B.

**Figure 2:**
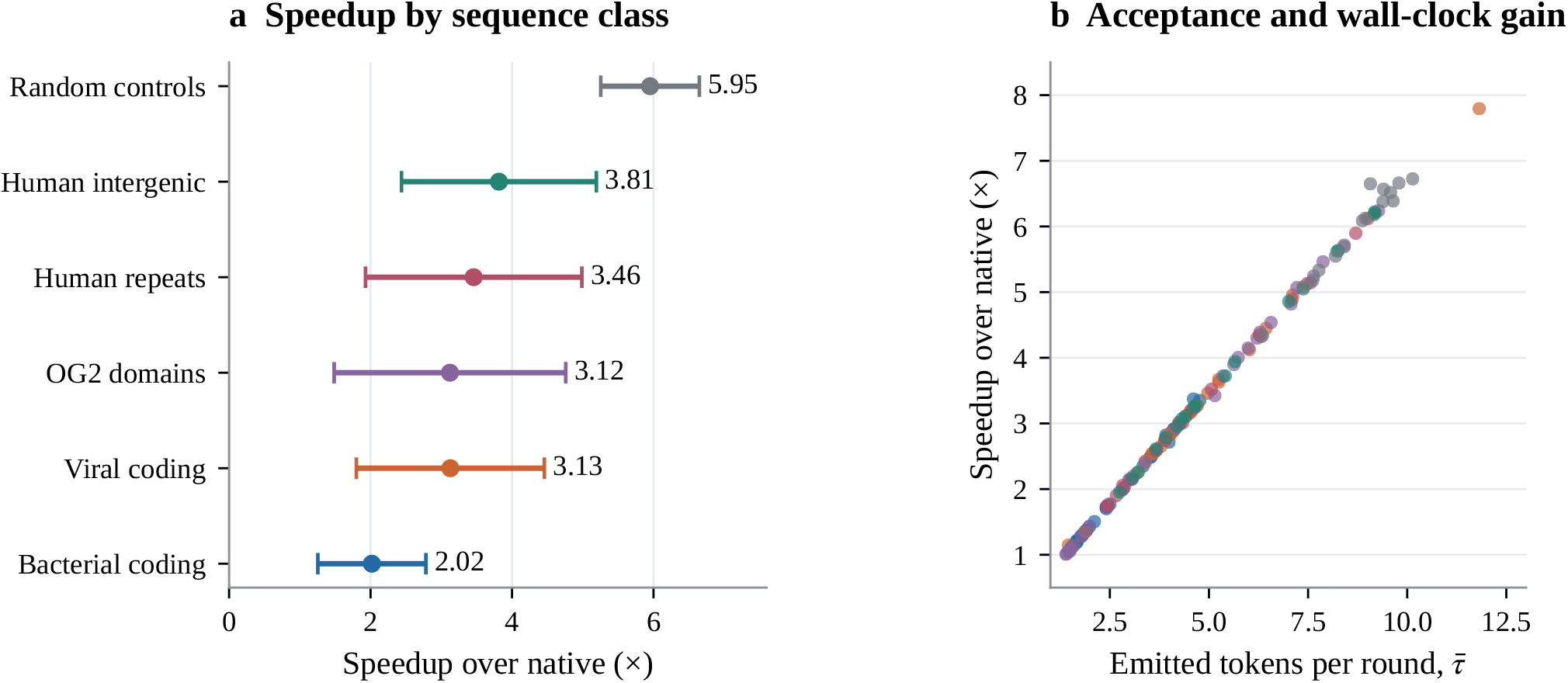
Evo2 7B deployment performance (48 prompts, three training seeds, rep-0). **(a)** Region means and population standard deviations across prompt–seed runs. **(b)** Speedup against mean emitted tokens per round for all 144 runs, colored by region as in (a). Higher acceptance generally increases speedup under the shared per-round latency budget.

The approximately linear relation between 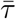 and speedup follows the cost decomposition in Equation 1. In the 7B loop, a draft costs about 4.3–4.5 ms and block verification about 22 ms. The target therefore processes several emitted tokens per expensive forward pass. The next two experiments test whether this balance survives larger targets and alternative drafter designs.

### 4.3 Acceleration extends to 20B and 40B targets

We retrain the same width-1024 drafter with *W* =64 and *γ*=12 for Evo2 20B and 40B. Hidden states are captured at layer 20 of 24 and layer 45 of 50, respectively, preserving approximately the relative depth of 7B’s layer 27. We retain the training objective and intended source mixture, with the architecture and draft length fixed across the larger targets.

The recipe yields 2.18*×* –2.46*×* on real sequences and 2.51*×* –2.78*×* on the full suite (Table 3). Increasing the budget from 10M to 30M changes the 20B all-48 mean by less than 0.5%; the three 40B budgets span less than 0.06*×* (Figure 3a). Within these single-training-seed runs, budget sensitivity remains low, consistent with the pattern observed at 7B. The broad region pattern also persists: random controls are easiest and bacterial coding is hardest.

**Table 3.** Larger-target decode speedups on one H20 (SDPA, one training seed, two decode repetitions per prompt). Each target has its own native reference.

| Target | Supervised positions | Real-43 | All-48 | $\bar{\tau}$ |
| --- | --- | --- | --- | --- |
| 20B | 10M | 2.18× | 2.53× | 4.07 |
| 20B | 30M | 2.22× | 2.51× | 4.03 |
| 40B | 10M | 2.44× | 2.72× | 4.03 |
| 40B | 30M | 2.46× | 2.78× | 4.08 |
| 40B | 80M | 2.42× | 2.72× | 4.00 |

**Figure 3:**
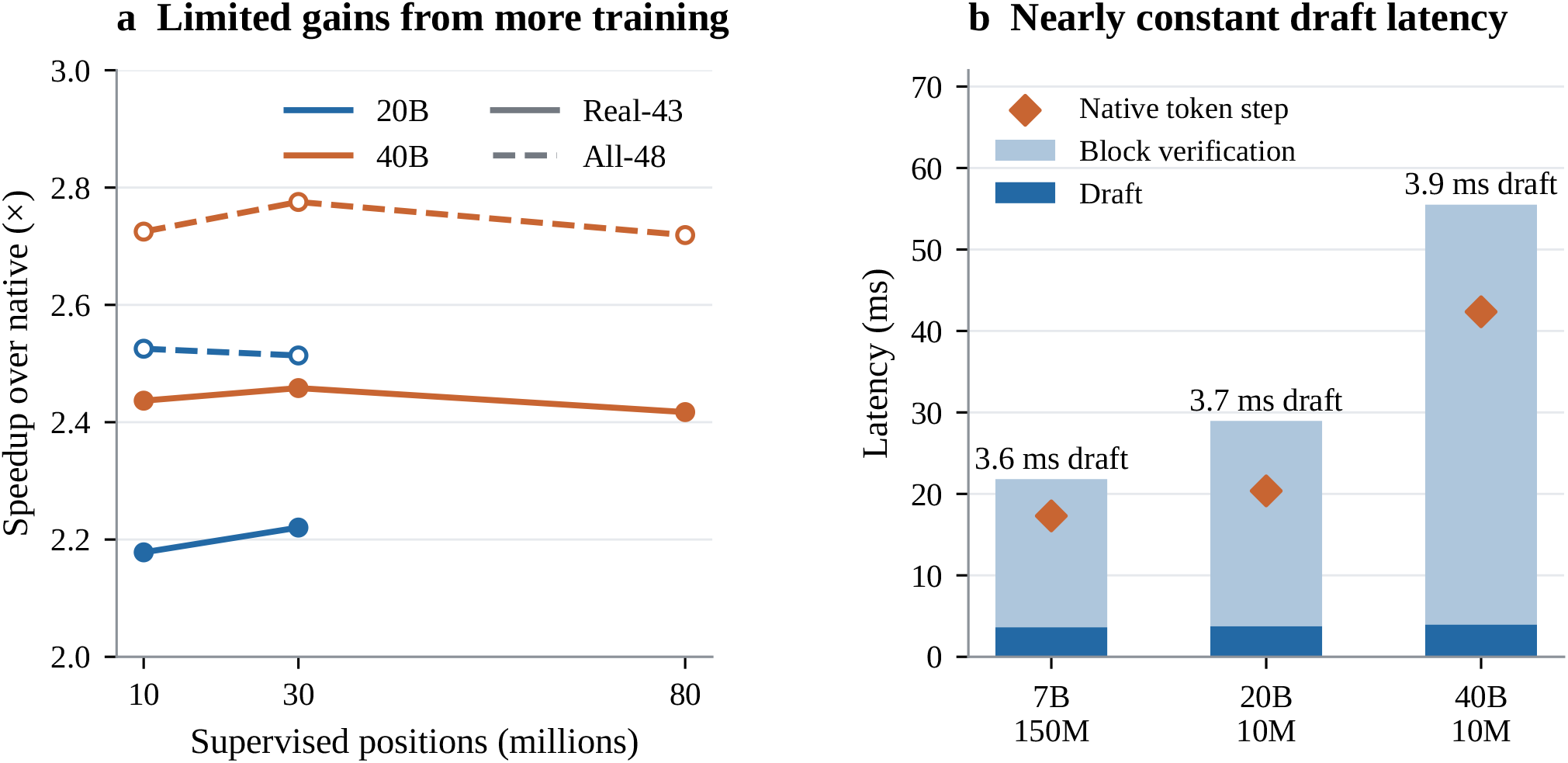
Transfer across target sizes on H20. **(a)** Budget curves for 20B and 40B; solid lines show real-43, dashed lines all-48. Each point uses one training seed and two decode repetitions. **(b)** Mean draft and block latencies measured inside the loop, with native per-token latency overlaid. The 7B checkpoint uses 150M training positions and flash attention; 20B/40B use 10M and SDPA. Bars show these two components, not total round latency.

Draft latency remains 3.6–4.0 ms across targets (Figure 3b). At similar acceptance lengths, its relative cost is lower for 40B than for 20B: 9.3% versus 18.4% of a native token step. This explains the higher 40B speedup without invoking better acceptance. A 7B retest on H20 yields 2.86*×* /3.18*×* (real/all), similar to the 4090 result. Its larger training budget and different attention backend preclude interpreting the three-model ordering as a pure scaling law. Mean emitted length is also lower for 20B/40B 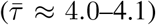 than for the H20 7B retest (4.50). Optimizing drafter capacity for each target size remains unexplored (Section 6).

Offline training takes 22–70 minutes for the 10M/30M cells and 3.0 hours for the 40B 80M cell on one H20. Training memory is about 20.5 GB because the teacher is absent from the optimization loop. Source-level accounting finds one reused pool: at 40B/80M, hg38 sampling corresponds to approximately 1.50 passes over the available hg38 window pool. The 20B/10M run receives no ncRNA windows despite their planned allocation; all nine sources are represented at 30M. The realized training mixture therefore differs in these two cells.

### 4.4 Draft latency governs speedup

We compare the parallel neural drafter with two autoregressive alternatives: First-*N* uses the target’s first 8 or 16 blocks, final normalization, and unembedding; an independent drafter uses the released Evo2 1B model. All arms use the same 7B target and the 48-prompt suite, with two decode repetitions at *γ* ∈ {2, 4, 7, 12} (Figure 4; exact values in Table 5).

**Figure 4:**
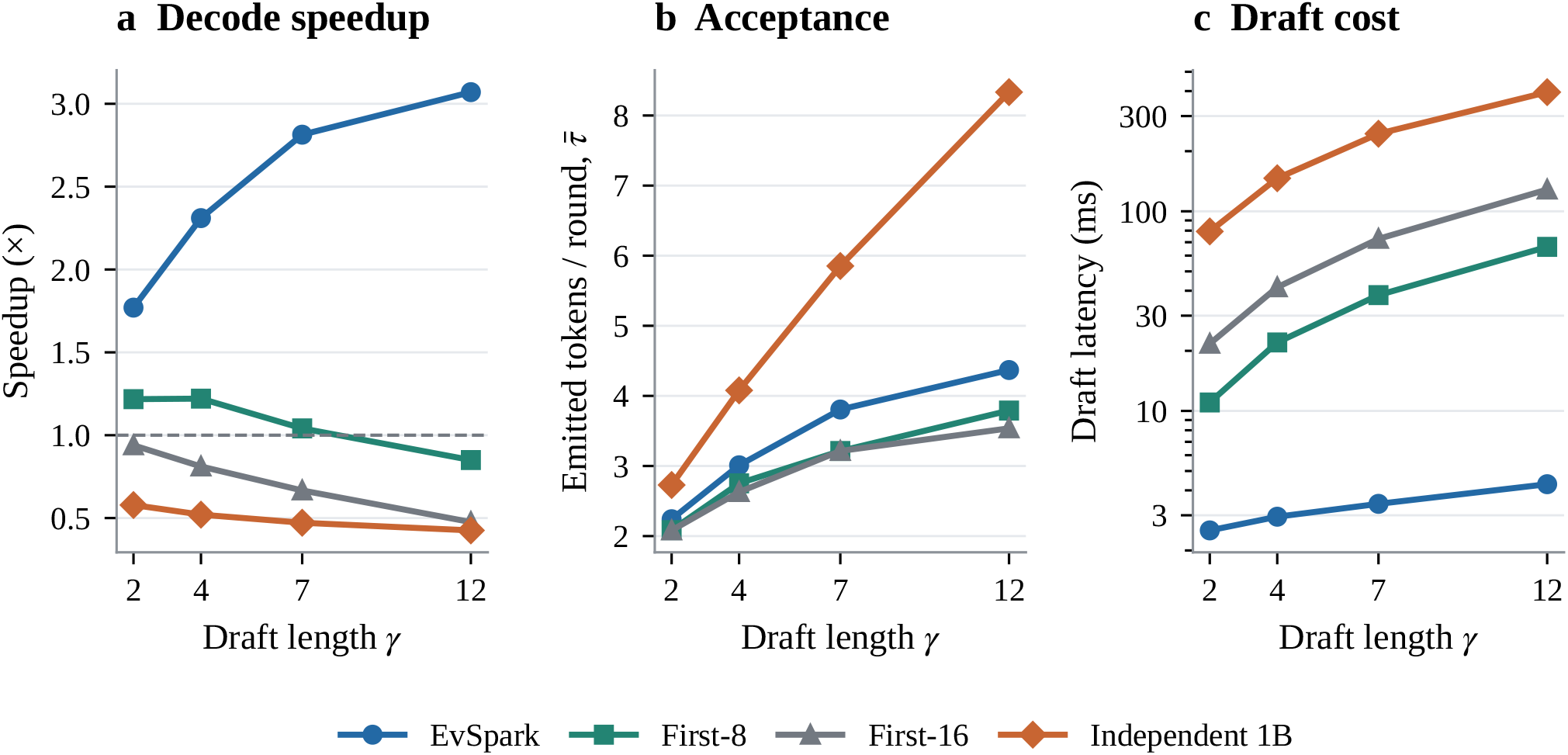
Drafter comparison on RTX 4090 (all-48, two decode repetitions). **(a)** Decode speedup, with native speed marked at 1*×*. **(b)** Mean emitted tokens per round. **(c)** Draft latency on a logarithmic scale. The independent 1B drafter has the highest acceptance at *γ*=12 but also the highest serial draft cost. These results characterize the evaluated implementations; the 1B arm uses unfused Hyena kernels and has an 8k context limit.

First-8 reaches 1.22*×* at *γ*=2 and 4, then falls below native speed at *γ*=12; First-16 is below native speed throughout. The 1B arm achieves 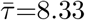 at *γ*=12, compared with 4.37 for EvSpark, but takes 395 ms to draft a block and achieves 0.43*×*. EvSpark’s draft cost is 2.5–4.3 ms over the tested lengths and its speedup rises from 1.77*×* to 3.07*×*. It is faster than these alternatives at every tested length. Serial autoregressive drafting is the latency bottleneck for the 1B arm in this 4090 implementation.

#### Prompt-lookup reference

A separate prompt-lookup baseline gives 1.276*×* and 1.282*×* for match lengths *k*=2 and 3 on all-48 (three sampling seeds). Here *k* is the lookup match length, not the number of drafted tokens *γ*. These measurements provide an additional training-free reference under our protocol. BioSpecDec reports 1.2–1.4*×* for small-model and truncated-target drafters on different targets [7]; its protocol and drafter classes differ from this prompt-lookup reference.

### 4.5 Batch scaling and multiple draft candidates

Batching independent sequences and proposing multiple candidates for one sequence spend parallel compute in different ways. We measure both, using *B* for independent sequences and *C* for candidates, to separate available forward-pass capacity from realized speculative speedup.

#### Independent-sequence batch measurements

On H20 with SDPA at 1k context, the 7B native step rises only from 21.1 to 22.3 ms between *B*=1 and 16. A 12-token parallel chunk stays near 26–27 ms through *B*=8, then increases to 41.8 ms at *B*=16 (Figure 5a,b). Memory limits are target-dependent: the first out-of-memory configurations are *B*=32 for 7B, *B*=8 for 20B, and *B*=2 for 40B in this 96 GB implementation. The corresponding 4090 probe shows a 5.1% increase in 12-token chunk time at *B*=4 and exhausts 48 GB at *B*=16.

**Figure 5:**
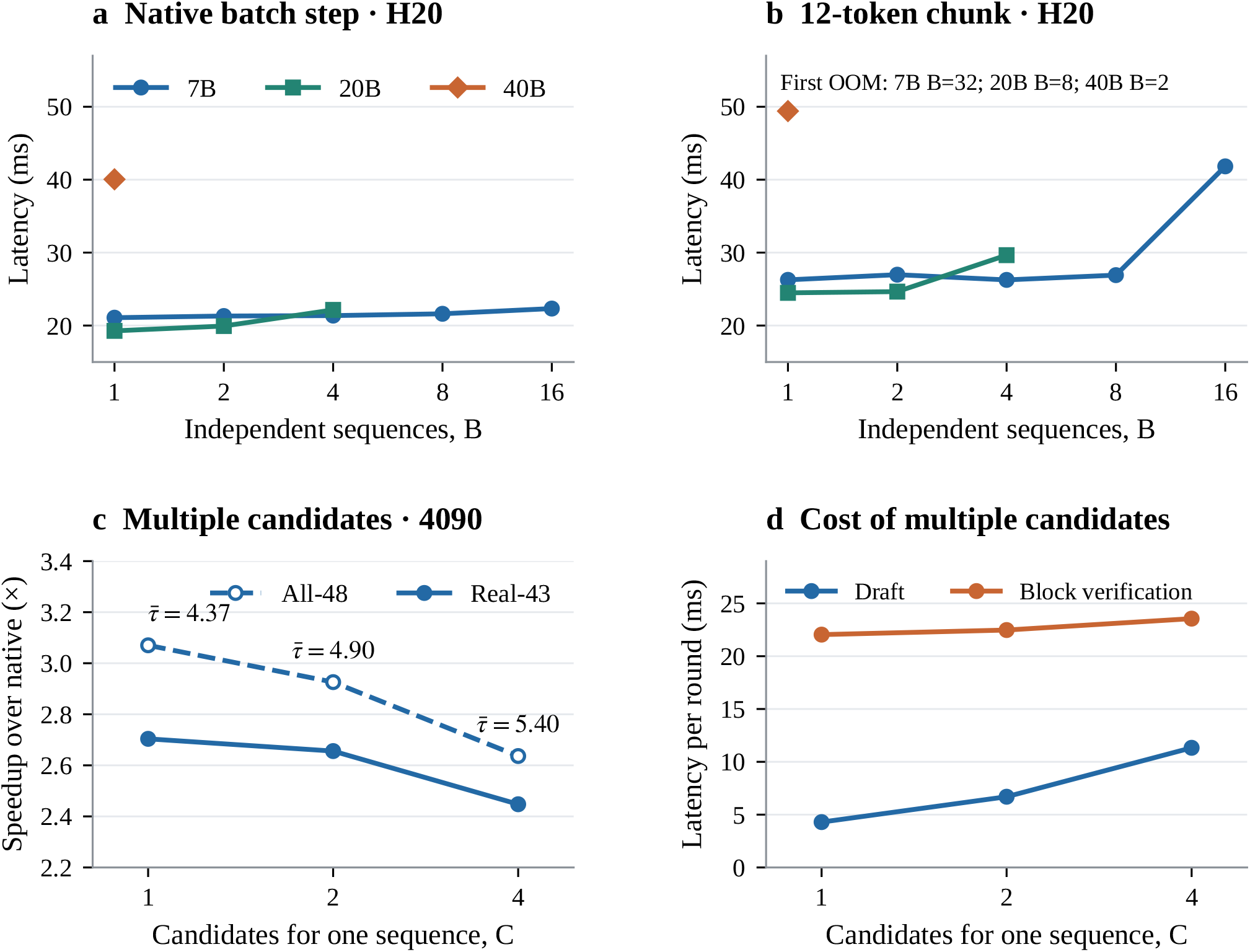
Batch capacity and candidate expansion are distinct measurements. **(a**,**b)** Native token-step and 12-token chunk latency versus independent sequence count *B* on H20, 1k context, SDPA; points are medians of repeated timed forwards. Curves end at the largest successful batch. **(c**,**d)** End-to-end multi-candidate decoding on 4090, *γ*=12, 48 prompts and two repetitions. Acceptance increases with candidate count *C*, but serial candidate sampling increases draft cost and lowers speedup.

These are forward-pass microbenchmarks. The chunk probe clears Hyena states to exercise the parallel path while retaining the specified attention context; it does not include accepted-prefix slicing, drafting, or scheduling of independently accepted lengths. The measurements quantify forward-pass capacity available for batch scheduling.

#### More candidates raise acceptance but slow decoding

We verify *C* candidate paths along the batch dimension and apply prefix-tree rejection sampling [25]. The winning accepted prefix selects the state and hidden-context buffer retained for the next round. With *C*=1, 2, 4, mean emitted length increases from 4.37 to 4.90 to 5.40, while all-48 speedup decreases from 3.07*×* to 2.93*×* to 2.64*×* (real-43: 2.70*×*, 2.66*×*, 2.45*×*). The shared trunk is computed once, but candidate sampling remains serial: draft time rises from 4.3 to 11.3 ms, while block verification rises only from 22.0 to 23.6 ms (Figure 5c,d). The additional accepted tokens do not offset that cost. We therefore retain a single candidate for the main performance experiments; parallelizing candidate sampling remains an optimization target.

### 4.6 Acceleration persists at long context and through 32k generation

Acceleration persists as attention-cache traffic increases (Figure 6). Held-out GTDB contigs yield seed-level mean speedups of 2.7–3.8*×* through their 51k context limit. On *E. coli* and *B. subtilis*, 262k-context speedups are 1.84–2.43*×* across five genome–seed combinations. Native throughput falls from approximately 43 to 25 tok/s between short and 262k context; EvSpark is also affected by these KV reads, but retains an advantage.

**Figure 6:**
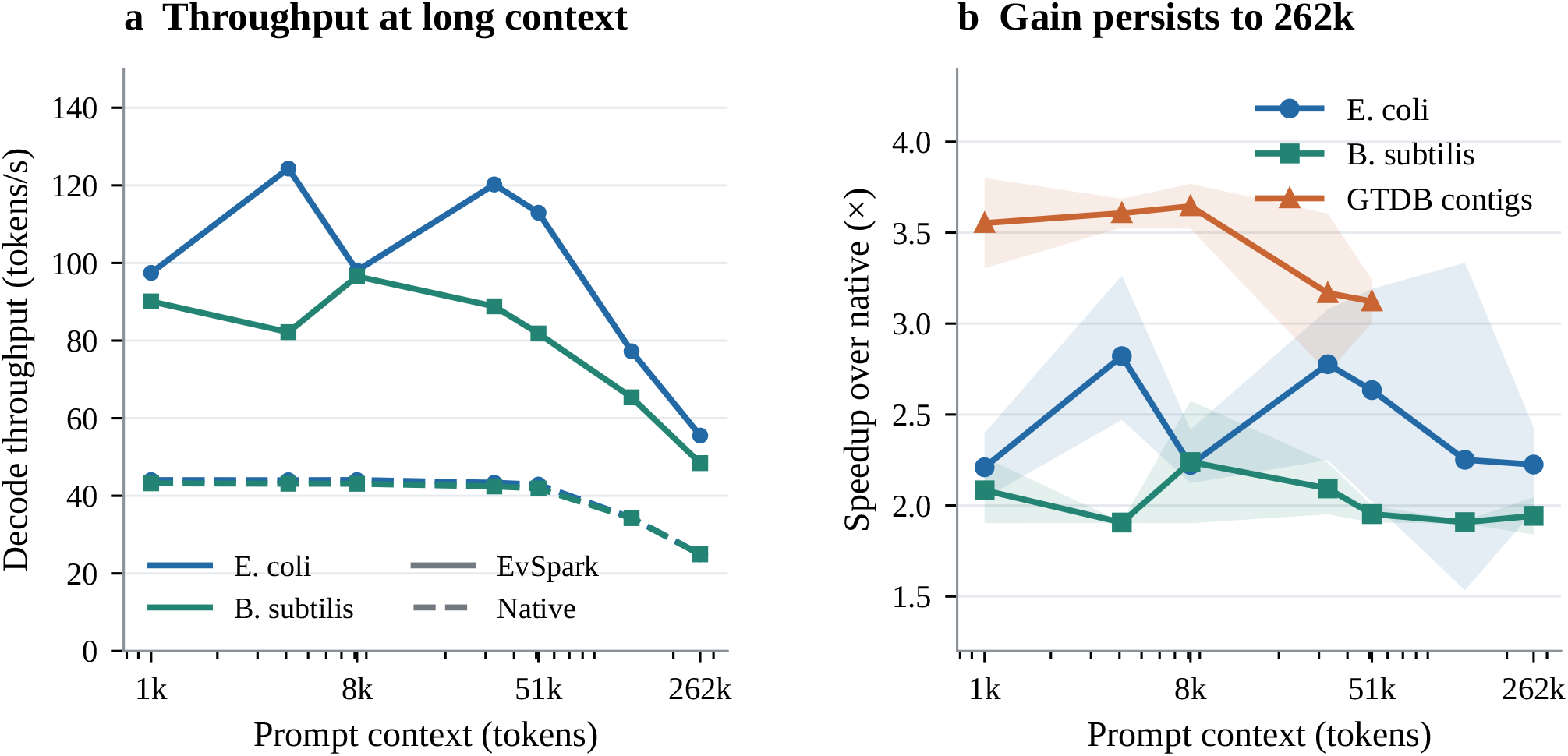
Context scaling with the 7B deployment drafter. **(a)** Mean native and speculative throughput on two bacterial genomes. **(b)** Speedup with bands spanning training-seed means: three seeds for *E. coli*, two for *B. subtilis* and GTDB. Genome-derived context points use two decode repetitions; GTDB uses three contigs and two repetitions per checkpoint. Both arms exclude the same chunked prefill.

Sustained generation from 1k prompts gives a complementary test. Across 32,768 generated tokens, four prompts and three decode repetitions, blockwise speedup shows no consistent downward trend. After discarding the first 1024-token block, repetition-level blockwise means range from 2.34 to 5.68*×*, depending on prompt and trajectory (Figure 10). The drafter’s fixed feature window and Hyena’s fixed-size state keep these components bounded; the target attention cost still grows with the generated sequence. Speculative block times sum recorded per-round draft, block-forward, verification and recovery components. The three repetitions use one deployment checkpoint.

### 4.7 Drafter configuration and diminishing returns from training

At a fixed 30M-position budget, increasing training and decode length from *γ*=4 to 12 raises all-48 speedup from 2.32*×* to 3.15*×*. Lengths 16 and 20 give 3.03*×* ; the per-seed ranges overlap the *γ*=12 range (Figure 7a). For a single architecture trained at *γ*=16, increasing decode length *γ*^*′*^ from 4 to 16 raises the two-seed mean from 2.21*×* to 3.14*×* (Figure 7b). This supports tuning decode length without retraining, while the separate training-length sweep also reflects changes in the learned drafter.

**Figure 7:**
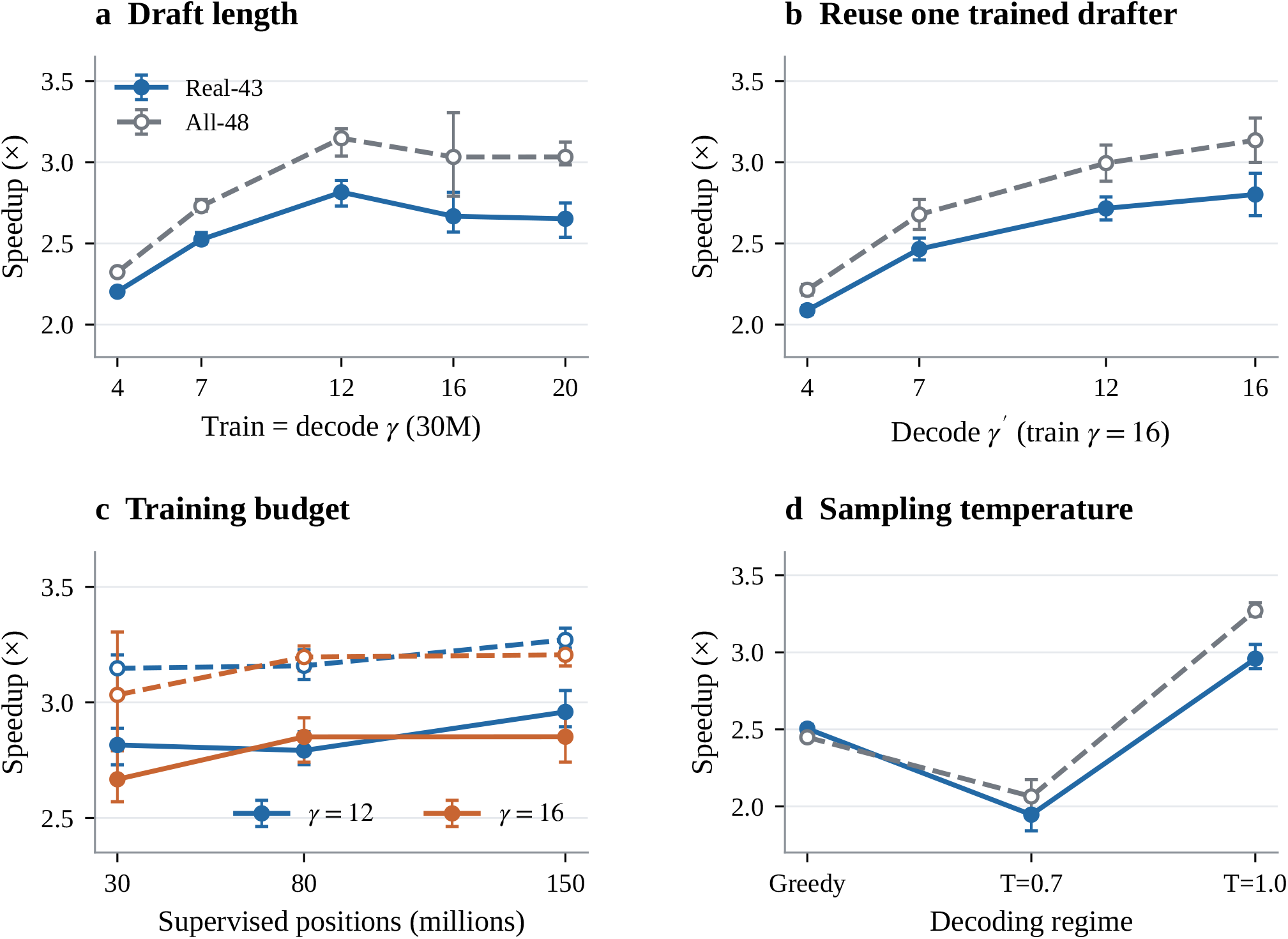
7B configuration effects. Solid lines: real-43; dashed lines: all-48. Error bars span training-seed suite means, not confidence intervals. **(a)** Training and decode length at 30M positions (three seeds). **(b)** Decode length of a drafter trained at *γ*=16 (two seeds). **(c)** Training budget for *γ*=12 and 16 (three seeds). **(d)** Decoding regime: greedy and *T* =0.7 pool decode repetitions for two seeds; *T* =1.0 uses the three-seed rep-0 grid.

Budget increases produce smaller changes. For *γ*=12, all-48 means are 3.15, 3.16, and 3.27*×* at 30M, 80M, and 150M positions; corresponding real-43 means are 2.82, 2.79, and 2.96*×*. For *γ*=16, real-43 is 2.85*×* at both 80M and 150M. Budget-related gains are comparable to the observed seed and trajectory variation. We use the configuration with the highest measured mean for deployment; the 30M setting offers a lower training cost.

The sampling regime also matters. The deployment drafter reaches about 2.45*×* under greedy decoding and 1.96–2.17*×* at *T* =0.7 (all-48, two seeds), below the *T* =1.0 configuration used for distillation. A lower temperature therefore need not increase speculative acceptance.

#### Features and data coverage

Matched-budget layer comparisons at *γ*=7 span 2.57–2.82*×* (30M positions, two seeds, 24-prompt protocol; Table 8). Final attention injection reaches 2.82*×* versus 2.65*×* for L27, but their ordering reverses at *γ*=12 (2.79 versus 3.03*×*). Layer rankings therefore depend on draft length and training budget; final attention also ranked lower at 0.5M positions. Width comparisons at 0.5M positions with one training seed give 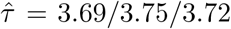 for *d* = 512*/*1024*/*2048. We select *d* = 1024 from these similar training-acceptance estimates; end-to-end capacity tuning is discussed in Section 6. Zeroing the Markov bias at evaluation has no consistent speedup benefit across feature layers (L27: 2.65 to 2.76*×*; final attention: 2.82 to 2.66*×*), and full-context feature injection does not outperform *W* = 64.

In a matched 30M-position, two-seed viral ablation, removing IMG/VR lowers held-out viral coding speedup from 2.23 to 1.77*×* (Figure 11). This provides direct evidence that source coverage can matter even when increasing total training positions gives little gain. These layer-injection and viral-source ablations retain their original 24-prompt, two-training-seed protocol and are not pooled with the main 48-prompt grid or the single-seed width proxy.

#### Training time

The *γ*=12 drafter trained at 30M positions takes 32.2 min on three RTX 4090s (mean of three seeds) and reaches 2.82/3.15*×* (real/all). The 150M deployment model reaches 2.96/3.27*×*; seeds 1/2 average 381.5 min on one GPU, while seed 3 takes 183.4 min on three GPUs. Table 7 reports all 33 training runs grouped by GPU count. These recorded training-loop times include evaluation, data handling and checkpointing, but exclude initialization and teacher-cache collection. Supplementary Data 3 gives the individual records. Teacher-data collection must be included when estimating the cost of training for a new target.

### 4.8 Decoding fidelity and small changes in sequence statistics

We compare EvSpark with native Evo2 decoding using token- and sequence-level tests.

#### Greedy and sampling checks

Across 48 prompts, six checkpoints, and 1024 generated tokens per test, greedy checks report zero non-tie divergences. They also identify 16–20 near-tie cases per checkpoint, documented with per-case logit evidence. In sampling tests with 24 prompts and two deployment checkpoints, 16 speculative chains are compared with an equal-size native reference. Two disjoint sets of 16 native chains provide a matched-*N* native–native KL reference. Speculative–native unigram KL is at or below that reference for 12/24 and 13/24 prompts in the two checkpoints (Figure 8a). The largest unigram TVD is 8.2% in repetitive sequence; non-repeat prompts remain below 4%.

**Figure 8:**
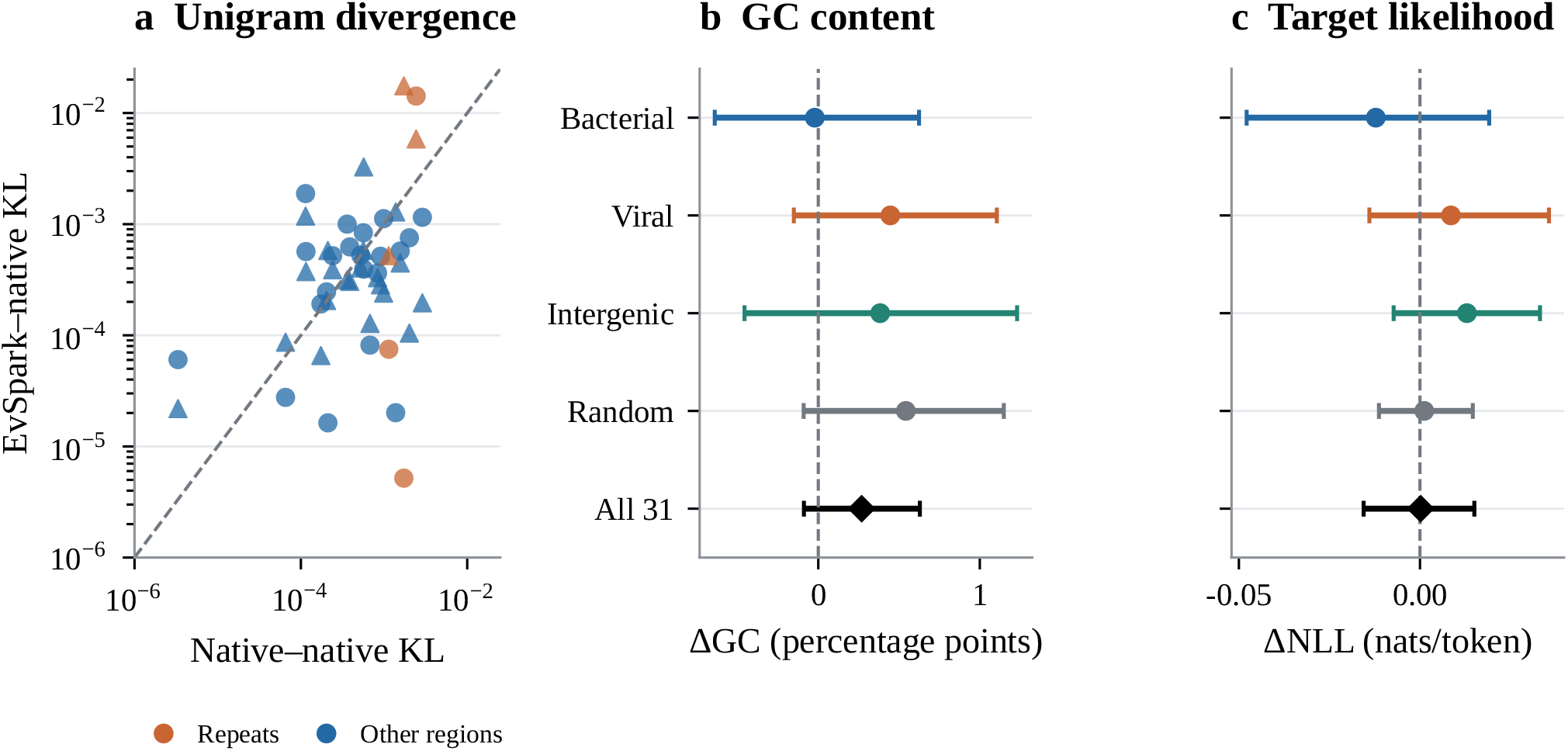
Sampling and sequence-statistic comparisons. **(a)** Per-prompt unigram KL against a matched-size native– native reference; circles and triangles denote deployment checkpoints trained with seeds 1 and 2, respectively. Orange marks repeat regions, and the dashed line is equality. **(b**,**c)** Exploratory sequence-level differences (EvSpark minus native) in GC content and target NLL. Points average prompts within each group; bars are 90% prompt-bootstrap intervals. The sequence experiment uses 31 prompts and 808 continuations per arm, separately from the 24-prompt KL experiment.

#### Sequence-level statistics

A separate experiment generates 808 continuations per arm across 31 prompts: 17 chains per bacterial prompt, 25 per viral prompt, 34 per intergenic prompt, and 40 per random prompt, all of length 1024. Averaging within-prompt differences gives ΔGC = +0.27 percentage points (90% prompt-bootstrap CI [ *−* 0.09, +0.63]), ΔNLL = +0.0002 nats/token ([*−* 0.016, +0.015]), and ΔORF = +4.4 bp ([*−* 9.5, +18.0]). These estimates show small average differences in the assessed statistics. Sampling and bootstrap details appear in Appendix D.

For all 31 prompts, 4-mer TVD between the two full chain pools is below the native split-half reference (means 0.063 and 0.087). The native halves contain fewer chains, increasing sampling variation in this descriptive reference (Appendix D).

### 4.9 Regulatory DNA design achieves a 1.57-fold workflow speedup

We asked whether faster decoding translates into shorter complete design workflows. We adapted Evo2’s computational chromatin-accessibility design task [1], starting from a 40,960 bp mouse genomic prefix and extending candidates in 128 bp blocks. Enformer [32] and four Flashzoi replicates [34] guided iterative selection of two branches; the original Borzoi mouse replicate-0 [33] was reserved for final checking. Two predefined accessibility patterns and two design seeds yielded four paired comparisons of 3,072 bp designs, with one final top-ranked sequence per backend and comparison. The search protocol and all individual results are given in Appendix E.

Separate full-workflow calibration selected native batch 8 and EvSpark batch 4 on the same devices. Including initial prefill, candidate generation, cache handling, scoring and final checking, mean design time decreased from 13.63 to 8.73 min. Paired speedups had a median of 1.57*×* and ranged from 1.42*×* to 1.70*×* (Figure 9c). All four outputs from each backend met the prespecified predictive qualification criterion: both search-ensemble and final-checker AUROC *≥* 0.90. Mean checker AUROCs were 0.975 for native and 0.977 for EvSpark. Each backend produced four exact-unique sequences; pooled 4-mer entropies were 7.81 and 7.67 bits, respectively. Within the common observed time window of 34.94 min, the prespecified design order yielded 4 qualified unique outputs with EvSpark and 2 with native.

**Figure 9:**
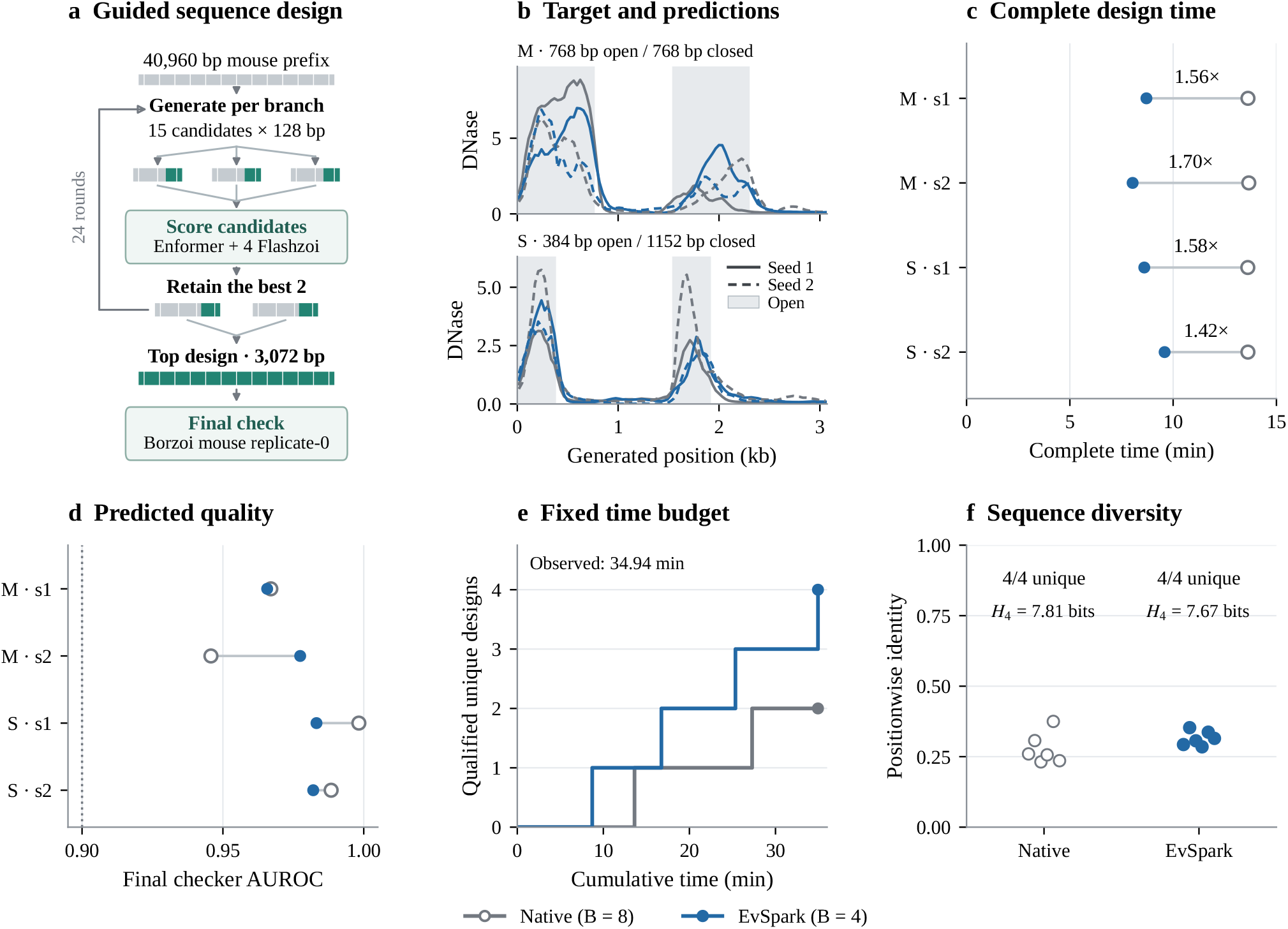
A computational regulatory-DNA design case study. **(a)** Fixed search workflow, shared by both backends; sequence strips are schematic, with green indicating new sequence. The first round starts with one branch. **(b)** Requested open intervals (gray shading) and final Borzoi DNase predictions for every output. Gray denotes native and blue EvSpark; solid lines denote design seed 1 (s1, 2026091901) and dashed lines seed 2 (s2, 2026091902). Each pattern shows all four outputs. M and S denote 768/768 bp and 384/1152 bp open/closed patterns, respectively. Raw prediction values are shown without display-specific renormalization, with a shared vertical scale within each pattern. The checker was excluded from search; no training-data independence is claimed. **(c)** Complete design times and paired speedups; rows identify the pattern and design seed. Model loading and warmup are reported separately. **(d)** Paired checker AUROCs in the same row order; the dotted line marks 0.90. Qualification additionally requires search-ensemble AUROC *≥* 0.90. **(e)** Cumulative exact-unique qualified final outputs in the fixed task order, restricted to the time window observed in both arms. **(f)** Pairwise positionwise sequence identity, exact-unique output counts and pooled 4-mer entropy (H4). Each arm contains four final sequences; pairwise identities are descriptive and are not independent design replicates.

**Figure 10:**
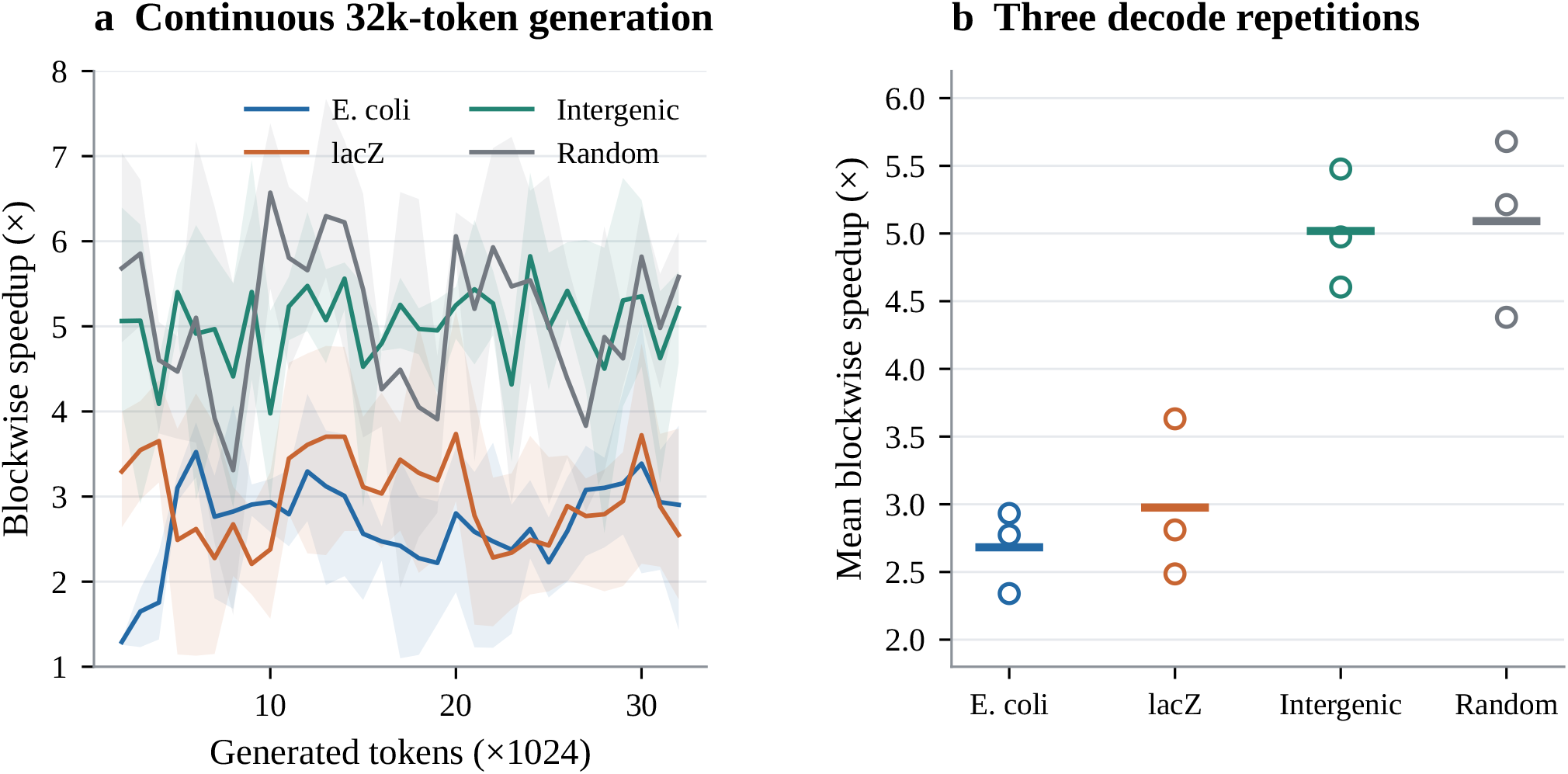
Sustained 7B generation from 1k-token prompts, one deployment checkpoint and three decode repetitions. **(a)** Blockwise speedup: lines show repetition means, bands their range. **(b)** Each point is a repetition’s mean over blocks 2–32; horizontal marks average the three repetitions. The first 1024-token block is discarded as warm-up. Speculative block times sum recorded per-round timing components; these are blockwise diagnostics rather than fullloop wall-clock ratios.

**Figure 11:**
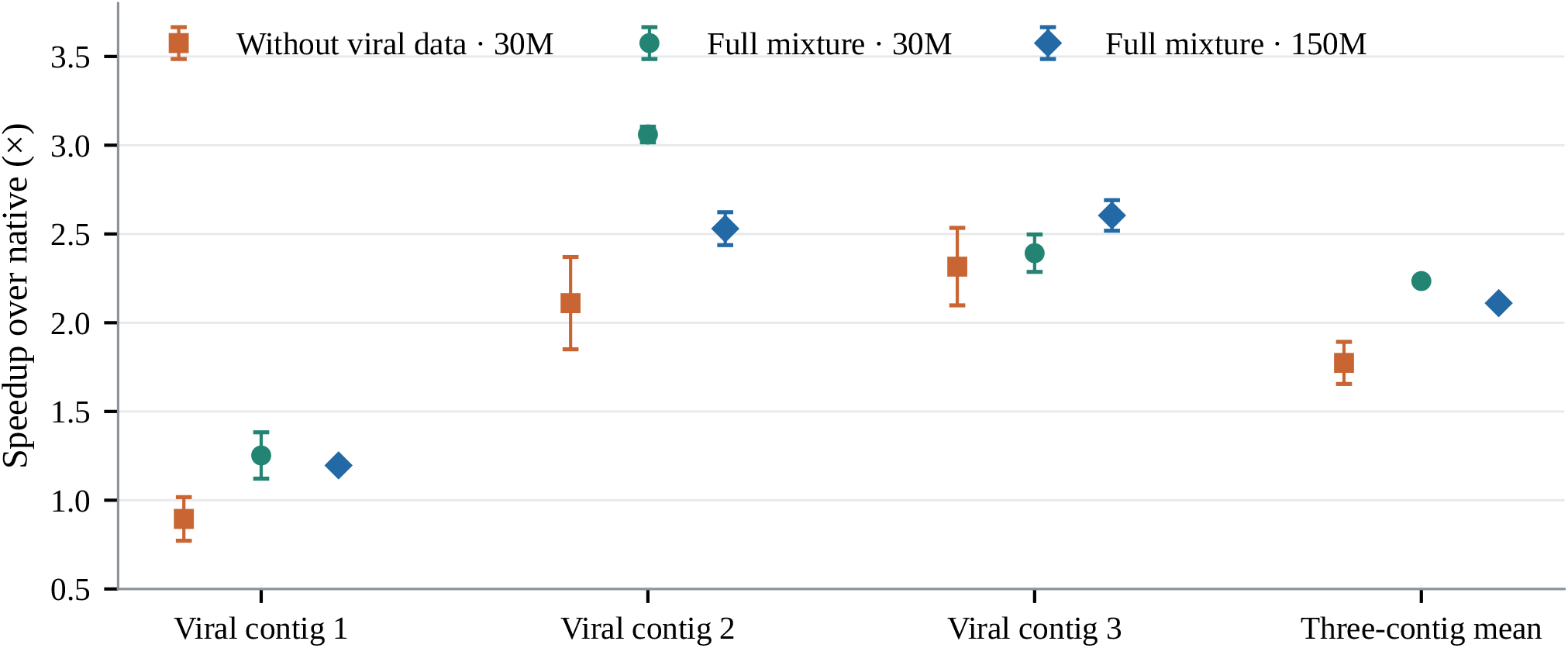
Viral-source ablation under one matched protocol. All three arms use *γ* = 7, the same three held-out viral prompts, 1024 generated tokens, and two training seeds from the original 24-prompt evaluation. Points show means across two training seeds and bars span their seed-level values. The final group averages the three prompts *within each seed* before computing its range.

**Figure 12:**
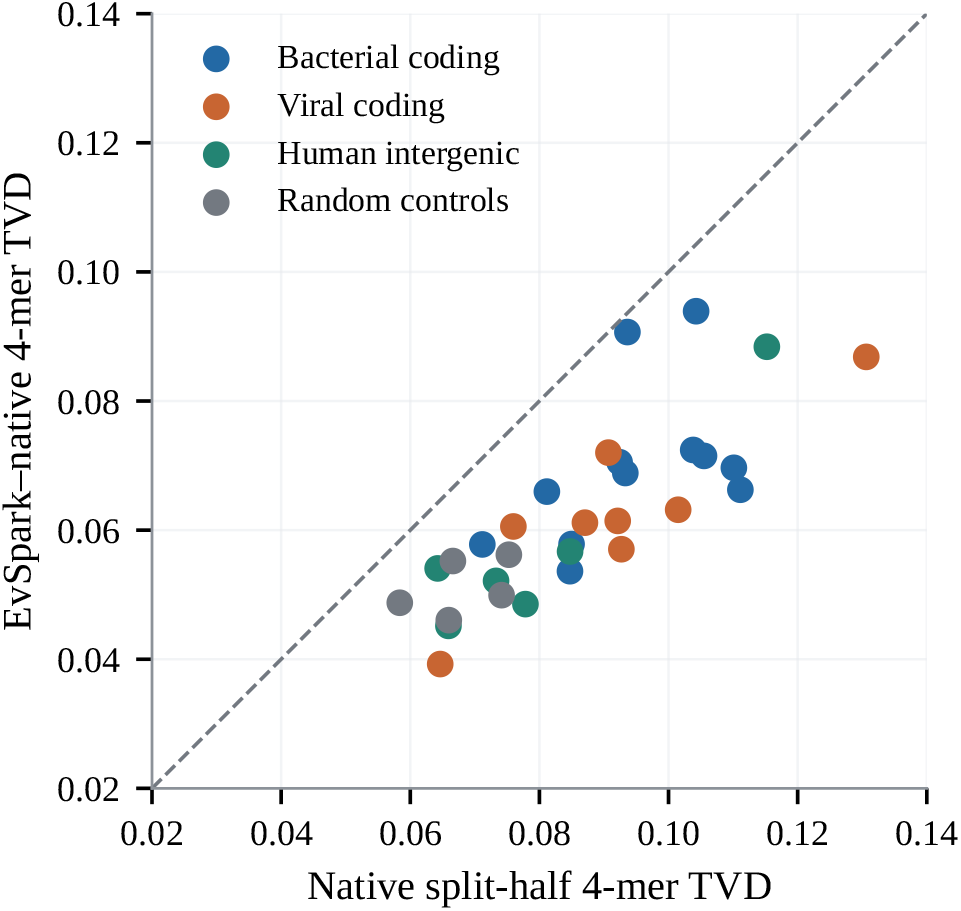
Per-prompt 4-mer TVD. The dashed line marks equal distances; the native split-half reference uses fewer samples than either full arm.

EvSpark reduces complete design time relative to a batched native baseline. Search scoring accounted for 47.9% of EvSpark’s mean complete-design time, showing how downstream costs limit the realized application speedup.

## 5 Related work

### Speculative decoding for biological sequence models

BioSpecDec [7] evaluates independent small-model and truncated self-drafters for Transformer-based genomic and protein language models, including DNAGPT [17], ProGen2 [18], and ProtGPT2 [19]. It reports average speedups of 1.2–1.4*×* and identifies Hyena integration and drafter distillation as future directions. EvSpark addresses those directions on Evo2’s hybrid architecture, with targets from 7B to 40B. Our First-*N* and 1B experiments instantiate comparable drafter classes on a shared 7B target. Differences in target, hardware, context, and measurement protocol prevent a direct comparison of headline speedups between papers.

### State-space and hybrid-model speculation

Snakes and Ladders [8], STree [9], and Mamba in the Llama [10] study speculative inference and state recovery for state-space or hybrid models. They establish that non-KV state need not preclude speculation. EvSpark’s contribution is specific to StripedHyena2: initial-state block verification and a joint slicing contract for finite convolution windows, modal IIR states, and attention caches. This complements work on Mamba [28] and Transformer–Mamba hybrids [29]; transferring the implementation to other recurrent operators would require their own intermediate-state construction and correctness checks.

### Learned and training-free drafters

Our parallel trunk, target-feature conditioning, and Markov bias follow DSpark [11]. We adapt its distillation pipeline to DNA and evaluate feature injection and training budget on a hybrid target, while leaving confidence-based scheduling unused. EAGLE [20], Medusa [21], Hydra [22], and EAGLE-2 [23] offer other feature-conditioned or multi-head drafting designs; DistillSpec [24] studies alignment through distillation. The general rejection-sampling framework comes from Leviathan et al. [14] and Chen et al. [15], following blockwise parallel decoding [16]. Prompt lookup [27], tree verification [25], and lookahead decoding [26] provide complementary ways to obtain or verify candidate continuations.

### Target architecture

Hyena’s long-convolution operators [13] and the StripedHyena family [12] supply the hybrid structure used by Evo2 [1]. EvSpark augments its native inference stack rather than requiring a different target architecture or retraining the target.

## 6 Limitations and future work

### Serving scope

The token-generation benchmarks concern single-sequence decode latency and exclude prefill. The regulatory-design case study additionally measures complete designs, including prefill, batched independent candidates and scoring, under a fixed search policy. Performance under arbitrary mixtures of concurrent requests remains to be evaluated. The earlier multi-candidate verifier produces one continuation and is distinct from these independent design candidates. Full-workflow gains depend on the costs of prefill, generation and downstream scoring.

### Hardware and model coverage

The 262k tests require the 48 GB 4090 variant. Larger targets are evaluated on one H20 with SDPA, and each larger-target budget uses one training seed. The measured acceleration spans these target sizes, with model-specific training budgets and attention backends. The slow 1B drafting path uses unfused Hyena kernels on Ada; optimized kernels or other hardware may change that comparison. We have not compared against all learned drafter architectures.

### Drafter capacity across target sizes

We retain the width-1024 drafter, context window, and training objective across target sizes, without a capacity or feature-layer search for 20B/40B. Their lower acceptance may reflect limitations of this transferred recipe, including drafter capacity, as well as differences in target distributions; distinguishing these factors requires matched capacity sweeps. Training budget and drafter capacity are separate optimization axes. More expensive target verification may justify a larger drafter, provided the relative gain in emitted length exceeds the relative increase in total round latency (Equation 1). Capacity and conditioning sweeps across training seeds should therefore assess end-to-end speedup alongside acceptance.

### Statistical and data scope

The 48-prompt suite samples a limited set of regions, with overlapping loci and unequal representation of biological diversity. Independent-locus controls support the broad profile but do not remove all dependence between prompts. Three training seeds cover the main 7B grid; decode trajectory variation can shift a suite mean by up to 0.44*×* across the measured repetitions. Older layer and source ablations use a separate 24-prompt protocol. The larger-target runs also have the realized-mixture exceptions disclosed in Section 4.3. Regulatory-design evaluation covers two patterns and two design seeds reused across patterns. Its quality and diversity summaries describe these eight outputs. Predictive qualification is assessed computationally; regulatory function has not been experimentally validated.

### Remaining performance limits

Coding sequences have lower acceptance across target sizes. Within the tested single-output verifiers, increasing training positions or the number of draft proposals does not reliably improve latency. Better coding-region proposals and scheduling across independent requests remain directions for improvement. Their value should be judged by end-to-end time, with training and data-collection costs included, rather than by acceptance alone.

## 7 Conclusion

EvSpark makes speculative decoding practical for Evo2 by combining a replay-free FIR/IIR/KV rollback protocol with a low-latency parallel drafter. The measured benefit extends from 7B to 20B and 40B targets and persists at long contexts. Drafter and candidate comparisons show that acceptance must be evaluated together with the cost of producing it. The resulting system also shortens a complete, predictor-guided regulatory-DNA design workflow against a batched native baseline.

### Code and reproducibility

Code is available at https://github.com/dhnihaoya/EvSpark, with released 7B, 20B and 40B drafters on Hugging Face and ModelScope. The numerical audit script recomputes reported benchmark aggregates from indexed JSON logs. The figure script additionally exports the plotted values and source-file hashes; figure captions specify which protocol each result uses. Supplementary Data 1 (JSON), 2 (FASTA) and 3 (CSV) contain evaluation prompts and source annotations, eight final design sequences, and 33 training-time records, respectively. These files, original benchmark and design records, and the audit and figure scripts are included in the accompanying source-data archive, indexed by file checksums.

## A. Exactness of the slicing protocol

The target and draft distributions below include the chosen temperature and top-*k* transform. All equalities in this appendix assume exact arithmetic. The invariant at a round boundary is that the target has consumed all emitted tokens *except the most recent one*, which is retained as the next anchor. This distinction is essential for both the indexing and the induction.

### Lemma 1

(Rejection-sampling correction). *For any fixed emitted prefix, let p*^*′*^ *and q*^*′*^ *be the corresponding target and proposal distributions. Draw x ∼ q*^*′*^, *accept it with probability* min(1, *p*^*′*^(*x*)*/q*^*′*^(*x*)), *and otherwise sample from the normalized positive residual* (*p*^*′*^ *− q*^*′*^)_+_. *The resulting token has distribution p*^*′*^.

*Proof*. The accepted mass at token *v* is min(*p*^*′*^(*v*), *q*^*′*^(*v*)). The total rejection probability is *Z* = ∑_*v*_ (*p*^*′*^(*v*) *− q*^*′*^(*v*))_+_. If *Z >* 0, residual sampling contributes (*p*^*′*^(*v*) *− q*^*′*^(*v*))_+_, so the total is *p*^*′*^(*v*). If *Z* = 0, the distributions coincide and every proposal is accepted. Proposal-zero events are never drawn. This is the standard speculative-sampling identity of [14, 15], applied conditionally at each output prefix. □

### Lemma 2

(Block-forward contract). *Given a legal state after L*_0_ *consumed tokens and a block z*_0_, …, *z*_*γ*_, *row i of the block-forward logits equals the sequential logits after consuming z*_0_, …, *z*_*i*_.

*Proof*. FIR convolution over the cached window and block evaluates the same filter at each position, with the same short-prefix convention. The parallel IIR computation evaluates the same linear recurrence, including the decayed initial state. Causal attention reads the same prefix keys and values. Positionwise feed-forward and normalization operations are unchanged. Composing these operators layer by layer gives the sequential logits at each position. The implementation checks this contract independently of the rejection sampler. □

### Lemma 3

(State restoration). *Let a round emit m tokens, possibly truncated by the remaining output budget. Selecting the retained state at block index j*^*\**^ = *m −* 1 *gives the sequential state immediately before the last emitted token. After full acceptance without budget truncation, the terminal block state already satisfies this property*.

*Proof*. The selected prefix consists of the old anchor and the first *m −* 1 emitted draft tokens. KV entries for this prefix are unchanged; its readable offset is *L*_0_ + *m*. The FIR slice selects the final *K −* 1 consumed filter inputs, or the native short-prefix state. The retained IIR state is the unique solution of the recurrence at *j*^*\**^. These are precisely the sequential states for that consumed prefix. For full acceptance, *m* = *γ* + 1, hence *j*^*\**^ = *γ* is the block’s terminal state. □

### Theorem 1

(Sequence-level exactness). *Under Lemmas 1–3, Algorithm 1 produces, for any fixed output budget, the same sequence distribution as sequential sampling from the transformed target. The target state and unconsumed anchor jointly satisfy the round-boundary invariant after every round*.

*Proof*. Prefill establishes the invariant. Within a round, Lemma 2 supplies the correct conditional target distributions. At each draft position, Lemma 1 gives the correct next-token law; a full-accept bonus is sampled directly from the corresponding target row. After a rejection or bonus, Lemma 3 restores the target to the state before the new anchor. The next round thus uses the correct conditional distributions after the emitted prefix. Induction over output positions gives the target law for any fixed-length output. Final-budget truncation simply retains its required prefix. With a deterministic argmax transform and the same tie-breaking rule, the same argument yields native greedy decoding. □

## B Detailed performance results

Table 4 gives the numerical region profile for Figure 2. Table 5 records the drafter sweep, and Table 6 separates context-length results by checkpoint. These experiments use different replication schemes; their suite means are not interchangeable estimates of one identical run.

**Table 4.** 7B deployment speedup by region. Values are pooled mean *±* population s.d. over three training seeds and *n* prompts per seed (rep-0). The s.d. describes heterogeneity, not uncertainty of the suite mean.

| Region | $n$/seed | Speedup (mean ± s.d.) |
| --- | --- | --- |
| Random (synthetic) | 5 | **5.95 ± 0.70** |
| Human intergenic (chr1 and chr21) | 6 | 3.81 ± 1.38 |
| Human repeat (hg38) | 6 | 3.46 ± 1.53 |
| Viral coding (IMG/VR valid) | 8 | 3.13 ± 1.33 |
| OG2 domain coverage (five sources) | 11 | 3.12 ± 1.64 |
| Bacterial coding | 12 | 2.02 ± 0.76 |
| **All-48** | 48 | **3.27 ± 1.68** |
| **Real-43** (primary) | 43 | **2.96 ± 1.47** |

**Table 5.** Autoregressive drafter comparison (all-48, 1024 tokens, two decode repetitions, RTX 4090). The neural row uses one deployment checkpoint; its 3.07× at *γ* = 12 is distinct from the three-seed rep-0 estimate of 3.27×.

| Drafter arm | $\gamma=2$ | $\gamma=4$ | $\gamma=7$ | $\gamma=12$ |
| --- | --- | --- | --- | --- |
| EvSpark (parallel) | 1.77× | 2.31× | 2.81× | **3.07×** |
| First-$N$ self-draft, $N=8$ | 1.22× | 1.22× | 1.04× | 0.85× |
| First-$N$ self-draft, $N=16$ | 0.94× | 0.81× | 0.67× | 0.48× |
| Independent 1B AR | 0.58× | 0.52× | 0.47× | 0.43× |

**Table 6.**
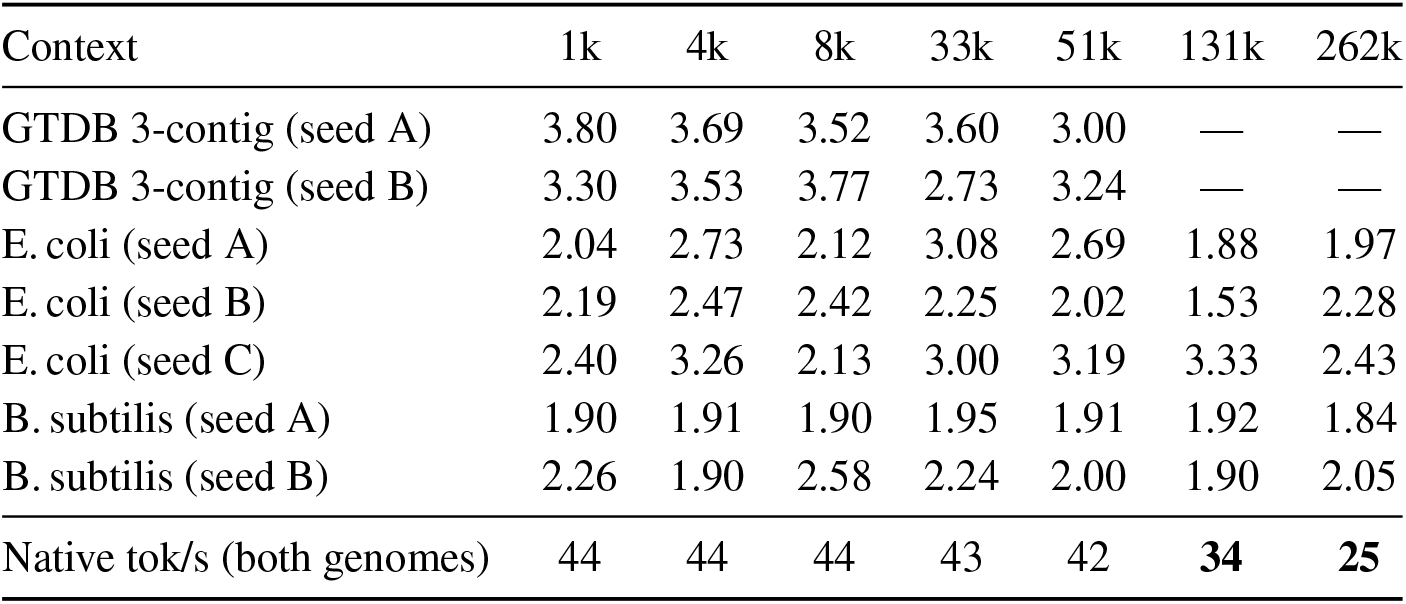
Context-length speedup by training seed. GTDB uses three contigs and two repetitions per checkpoint; the genome curves use two repetitions. Both arms use the same excluded 1k-chunked prefill. Values are speedups except for the final native-throughput row, which equally averages all seed–repetition runs across the two genomes.

## C Configuration and distillation details

Table 7 consolidates draft length, training budget, and recorded training time for the main grid. Layer and domain ablations retain their original 24-prompt, two-training-seed protocol (Table 8, Figure 11).

**Table 7.** 7B configuration performance and recorded training-loop wall-clock time. (a) Three training seeds, 48 prompts, rep-0; parentheses span seed-level all-48 means. (b) Time and step-external fraction grouped by actual GPU count; means (ranges) are shown, or a single value for one run. All GPUs are RTX 4090s.

| (a) Decoding performance |  |  |  |  |  |
| --- | --- | --- | --- | --- | --- |
| $\gamma$ | Positions (M) | Real-43 | All-48 (seed range) | $\bar{\tau}$ | |
| 4 | 30 | 2.20 | 2.32 (2.32–2.33) | 3.04 |  |
| 7 | 30 | 2.52 | 2.73 (2.70–2.77) | 3.71 |  |
| 7 | 80 | 2.68 | 2.88 (2.73–3.03) | 3.91 |  |
| 7 | 150 | 2.60 | 2.81 (2.78–2.84) | 3.83 |  |
| 12 | 30 | 2.82 | 3.15 (3.04–3.21) | 4.55 |  |
| 12 | 80 | 2.79 | 3.16 (3.10–3.23) | 4.56 |  |
| 12 | 150 | 2.96 | 3.27 (3.24–3.32) | 4.71 |  |
| 16 | 30 | 2.67 | 3.03 (2.79–3.30) | 4.59 |  |
| 16 | 80 | 2.85 | 3.20 (3.14–3.24) | 4.83 |  |
| 16 | 150 | 2.85 | 3.21 (3.16–3.26) | 4.85 |  |
| 20 | 30 | 2.65 | 3.03 (2.98–3.12) | 4.69 |  |

| (b) Training time |  |  |  |  |  |
| --- | --- | --- | --- | --- | --- |
| $\gamma$ | Positions (M) | GPUs | Seeds | Time (min) | Eval + other (%) |
| 4 | 30 | 3 | 1–3 | 96.0 (93.9–97.4) | 30.7 (26.5–33.7) |
| 7 | 30 | 3 | 1–3 | 52.0 (45.9–55.4) | 29.3 (19.2–35.1) |
| 7 | 80 | 3 | 1–3 | 151.8 (145.7–162.9) | 36.6 (33.3–42.8) |
| 7 | 150 | 3 | 1–3 | 274.1 (265.0–282.6) | 31.6 (27.3–35.4) |
| 12 | 30 | 3 | 1–3 | 32.2 (31.2–33.5) | 32.5 (27.9–35.6) |
| 12 | 80 | 1 | 1, 2 | 212.4 (211.7–213.1) | 42.1 (41.7–42.4) |
| 12 | 80 | 3 | 3 | 77.4 | 25.1 |
| 12 | 150 | 1 | 1, 2 | 381.5 (378.1–384.9) | 39.5 (39.2–39.7) |
| 12 | 150 | 3 | 3 | 183.4 | 22.7 |
| 16 | 30 | 1 | 1, 2 | 53.3 (47.8–58.8) | 33.1 (26.2–40.0) |
| 16 | 30 | 3 | 3 | 27.2 | 17.8 |
| 16 | 80 | 3 | 1–3 | 71.9 (66.9–75.6) | 17.9 (15.6–21.9) |
| 16 | 150 | 3 | 1–3 | 131.1 (116.5–139.4) | 19.3 (16.4–22.1) |
| 20 | 30 | 3 | 1–3 | 25.1 (23.6–26.8) | 15.7 (13.5–17.0) |

The interval includes training-loop evaluation, checkpointing and final checks, excluding initialization and teacher-cache collection. Eval + other is the step-external fraction, not evaluation-only time. Seeds 1–3 are 20260821–20260823. Supplementary Data 3 gives all 33 records. Execution modes were not a controlled comparison of training efficiency.

**Table 8.**
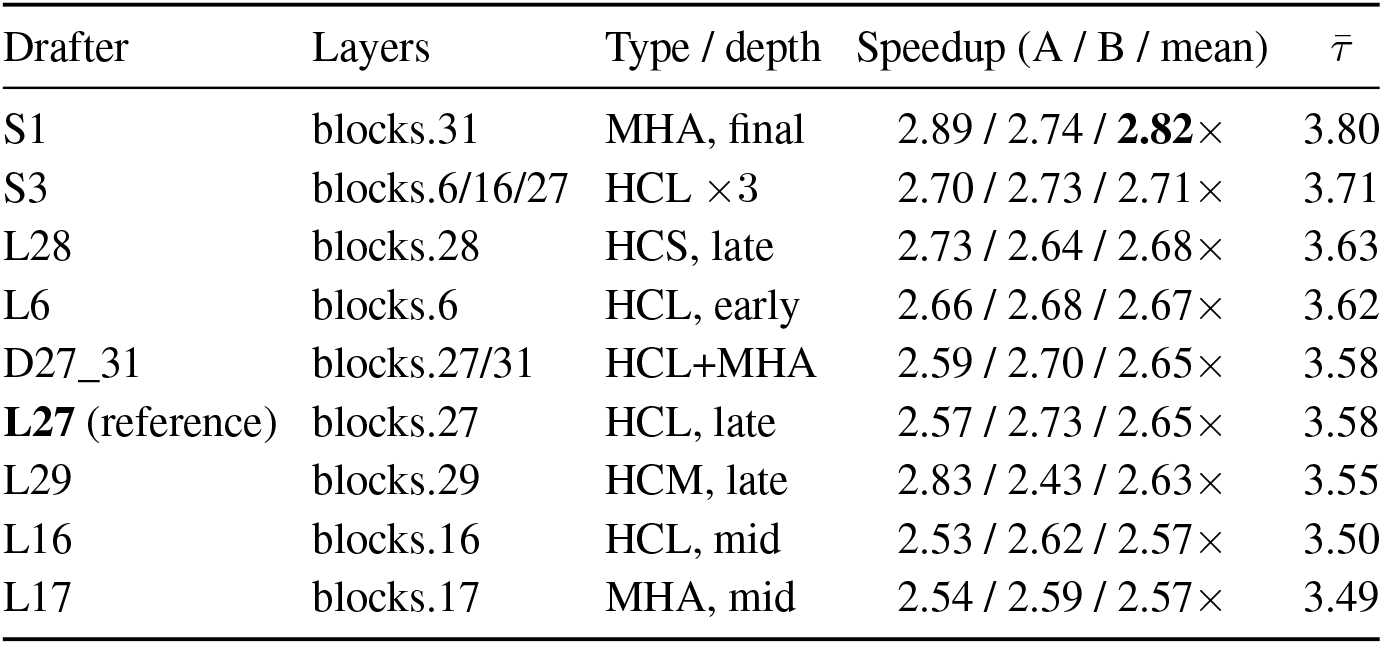
Matched-budget injection-layer comparison: 30M positions, *γ* = 7, two training seeds, 24-prompt protocol. Speedup lists the two seed means and their average; 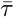 is the mean emitted length.

| Drafter | Layers | Type / depth | Speedup (A / B / mean) | $\bar{\tau}$ |
| --- | --- | --- | --- | --- |
| S1 | blocks.31 | MHA, final | 2.89 / 2.74 / **2.82**× | 3.80 |
| S3 | blocks.6/16/27 | HCL $\times 3$ | 2.70 / 2.73 / 2.71× | 3.71 |
| L28 | blocks.28 | HCS, late | 2.73 / 2.64 / 2.68× | 3.63 |
| L6 | blocks.6 | HCL, early | 2.66 / 2.68 / 2.67× | 3.62 |
| D27_31 | blocks.27/31 | HCL+MHA | 2.59 / 2.70 / 2.65× | 3.58 |
| **L27** (reference) | blocks.27 | HCL, late | 2.57 / 2.73 / 2.65× | 3.58 |
| L29 | blocks.29 | HCM, late | 2.83 / 2.43 / 2.63× | 3.55 |
| L16 | blocks.16 | HCL, mid | 2.53 / 2.62 / 2.57× | 3.50 |
| L17 | blocks.17 | MHA, mid | 2.54 / 2.59 / 2.57× | 3.49 |

### Cache, supervision and source allocation

The stored token IDs supply cross-entropy labels and reference predecessors. Top-eight teacher entries and residual mass support reconstruction of the top-four, temperature-1 targets. Int8 feature values are stored with their scales; late-layer scales use fp32 to avoid overflow. This codec applies to offline teacher data only. Target weights, KV cache and recurrent state are not quantized by the distillation codec.

Supervised-position budgets count draft prediction positions evaluated by the loss. With *B*_*a*_ anchors per rank and draft length *γ*, a synchronized step contributes *B*_*a*_*γ* positions per rank; the reported budget sums across ranks. It counts neither unique corpus bases nor distinct cached windows. Different anchors can supervise overlapping positions in a window. Windows are allocated in proportion to the intended source weights, subject to a cap of two passes over each source’s available windows per allocation epoch; unused allocation is redistributed. Reaching the supervision budget can stop an allocation before all planned windows are visited. An allocation epoch therefore does not imply that every source was visited equally or that every planned window was consumed. Table 9 lists intended weights; the realized 20B/10M ncRNA omission and 40B/80M hg38 revisit are reported in Section 4.3.

**Table 9.** Intended source-sampling mixture. Weights are independent of cache size; available-window caps and the stopping budget can alter realized exposure. Rounded percentages are shown; unrounded weights sum to one.

| Source | Target weight (%) |
| --- | --- |
| GTDB | 20 |
| mRNA | 31.5 |
| IMG/VR | 10 |
| Eukaryotic sequence | 10 |
| NCBI coding sequence | 9.5 |
| ncRNA | 5.0667 |
| Organelles | 2.5333 |
| Promoters | 1.9 |
| hg38 | 9.5 |

### Recorded wall-clock interval

Timing begins after model initialization and evaluation-pack preparation, and ends after final checkpointing and consistency checks. For each run, the elapsed time is read directly from its training record. The timed training-step component covers forward, loss, backward, gradient clipping, optimizer/scheduler updates and in-step synchronization; compilation on an initial timed step is not removed. *Eval + other* is computed per run as

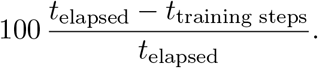

It includes evaluation, data preparation, logging, saving and final checks. Evaluation alone was not separately timed. Means and ranges are taken within groups sharing the same GPU count, without converting elapsed time into GPU-hours or combining single-GPU and distributed runs into one time mean.

## D Sequence-level statistical diagnostics

The frozen sequence-generation protocol contains 31 prompts and 808 continuations per arm, with unequal chain counts across regions (Table 10). Arms use independent random streams; all metrics use the 1024-token continuation, conditional on its prompt where required. GC is (*G* + *C*)*/*(*A* + *C* + *G* + *T* ); chains without canonical bases have no GC estimate. The longest complete ORF is scanned over all six frames, from ATG to an in-frame stop, with a 60-bp minimum and zero if none is found. Target NLL is the mean negative log probability under teacher-forced, full-vocabulary Evo2 logits. This likelihood metric is distinct from the top-*k* sampling law.

**Table 10.**
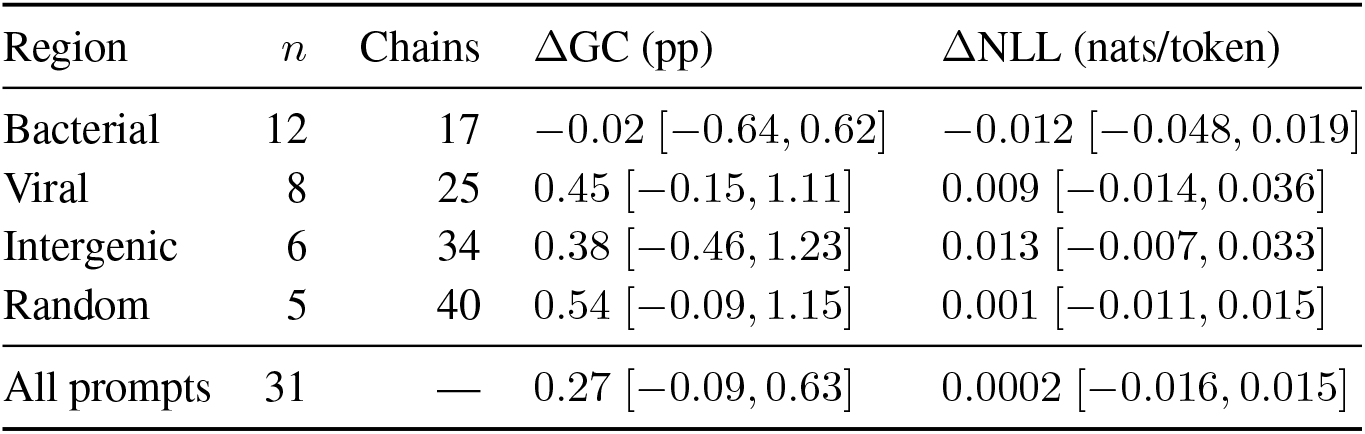
Exploratory sequence-statistic differences, EvSpark minus native. Intervals are 90% prompt-bootstrap intervals. *n* counts prompts; chains are per prompt *per arm*.

| Region | $n$ | Chains | $\Delta\text{GC}$ (pp) | $\Delta\text{NLL}$ (nats/token) |
| --- | --- | --- | --- | --- |
| Bacterial | 12 | 17 | $-0.02$ $[-0.64, 0.62]$ | $-0.012$ $[-0.048, 0.019]$ |
| Viral | 8 | 25 | $0.45$ $[-0.15, 1.11]$ | $0.009$ $[-0.014, 0.036]$ |
| Intergenic | 6 | 34 | $0.38$ $[-0.46, 1.23]$ | $0.013$ $[-0.007, 0.033]$ |
| Random | 5 | 40 | $0.54$ $[-0.09, 1.15]$ | $0.001$ $[-0.011, 0.015]$ |
| All prompts | 31 | — | $0.27$ $[-0.09, 0.63]$ | $0.0002$ $[-0.016, 0.015]$ |

### Original pooled analysis

The initial analysis pools chains within each region and resamples chains without prompt stratification, using 10,000 bootstrap resamples. Prespecified equivalence margins are 1.5 percentage points for GC, 50 bp for longest ORF, 0.03 nats/token for NLL, and 0.03 for 4-mer TVD. Signed mean differences use 90% percentile intervals, required to fall wholly inside their margins. For TVD, the upper endpoint of the two-sided 95% percentile interval must fall below its margin. The joint criterion is not met; margins and results are retained unchanged in Table 11. These study-specific diagnostic margins were not calibrated to downstream biological effects. The criterion assesses the selected sequence statistics under this analysis; it does not test the rejection-sampling identity. Resampling without prompt stratification allows prompt composition to vary, and its TVD bootstrap interval can lie above the pooled point estimate.

**Table 11.** Original pooled sequence-statistic analysis. EvSpark minus native for signed differences; TVD is unsigned. Brackets are 90% bootstrap intervals for differences and two-sided 95% intervals for TVD. Margins are *±*1.5 pp, *±*50 bp, *±*0.03 nats/token and TVD *<* 0.03.

| Region | $\Delta\text{GC}$ (pp) | $\Delta\text{ORF}$ (bp) | $\Delta\text{NLL}$ | 4-mer TVD |
| --- | --- | --- | --- | --- |
| Bacterial | $-0.02$ $[-1.17, 1.18]$ | $2.9$ $[-29.1, 35.0]$ | $-0.012$ $[-0.064, 0.040]$ | $0.021$ $[0.028, 0.045]$ |
| Viral | $0.38$ $[-0.94, 1.66]$ | $21.5$ $[-13.5, 56.5]$ | $0.009$ $[-0.057, 0.076]$ | $0.023$ $[0.029, 0.050]$ |
| Intergenic | $0.38$ $[-0.85, 1.62]$ | $-9.0$ $[-31.6, 12.9]$ | $0.013$ $[-0.019, 0.046]$ | $0.021$ $[0.027, 0.049]$ |
| Random | $0.54$ $[-0.15, 1.24]$ | $-3.0$ $[-25.2, 18.9]$ | $0.001$ $[-0.009, 0.012]$ | $0.023$ $[0.028, 0.039]$ |

### Prompt-level diagnostics

The subsequent exploratory analysis first takes the mean EvSpark-minus-native difference within each prompt and then weights prompts equally. Its 90% intervals use 10,000 bootstrap resamples of prompts, preserving the paired comparison between arms. Table 10 reports these prompt-level estimates. Four-mer TVD compares complete chain pools within each prompt; the native odd/even split uses fewer chains and supplies a descriptive reference, not matched-*N* calibration. This differs from the separate 24-prompt unigram KL experiment, which compares groups of 16 chains on both sides.

## E Regulatory-design case-study protocol

### Workload and search

We use Evo2 7B in bf16 with the layer-27, *γ*=12, 150M-position drafter (training seed 1), temperature 1 and top-*k*=4. The 40,960 bp prefix is mm39 chrX:[52,010,968,52,051,928), using zero-based, half-open coordinates. Scoring backgrounds retain upstream sequence ending at 52,051,928 and downstream sequence beginning at 52,123,468; the intervening reference segment is replaced by the generated design. At each round, each retained parent generates 15 independent 128 bp extensions. After scoring, the two candidates with the lowest loss are retained globally. The first round has one parent; 24 rounds evaluate 705 candidate extensions and return the highest-ranked 3,072 bp sequence. Prefix states are reused, with private candidate state and separate random streams. This candidate batch is distinct from the multiple-proposal, single-output verifier in Section 4.5.

The two targets alternate 768 bp open/768 bp closed (M) and 384 bp open/1,152 bp closed (S). Each is run with design seeds 2026091901 and 2026091902 under both backends, yielding eight completed designs. Calibration seed 2026091801 is separate from these eight outputs.

### Prediction coordinates and selection loss

All predictors use mouse DNase track ENCFF872IES. Enformer receives 196,608 bp including 40,960 bp upstream of the design; Flashzoi and the final Borzoi checker receive 524,288 bp including 163,840 bp upstream. The generated interval begins at the first predicted bin, with resolutions of 128 bp and 32 bp, respectively. Future positions are filled with the fixed downstream background and excluded from the loss and AUROC. Enformer uses fp32 with the TensorFlow-gamma implementation; Flashzoi uses fp16 autocast; the original Borzoi checker uses fp32.

For each candidate, let *e* be Enformer’s full 896-bin profile divided by its maximum, and let *f*_*r*_ be Flashzoi replicate *r* divided by the common maximum over all four 6,144-bin profiles. Define *f* = mean_*r*_(*f*_*r*_) *−* std_*r*_(*f*_*r*_), using the population standard deviation. For generated length *L*, selection minimizes

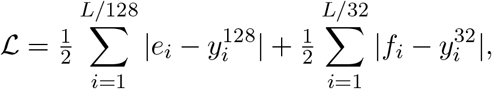

where *y*^*b*^ is the prescribed binary target at resolution *b*. Search-ensemble AUROC is the arithmetic mean of the AUROCs of *e* and *f* . The final checker is the original Borzoi mouse replicate-0, excluded from selection, evaluated on the 96 generated bins. Qualification requires both ensemble and checker AUROC *≥* 0.90. This is a predictive criterion; the checker’s training data are not claimed to be independent.

### Calibration and timing

Both arms use the same GPU pair: a 48 GB RTX 4090 for generation and a 24 GB RTX 4090 for scoring, with scoring batch 4. Full-workflow calibration tests native batch sizes 1/2/4/8 and EvSpark 1/2/4, selecting batch 8 and 4 solely by elapsed time (Table 12). Quality does not select the batch, and the unqualified native batch-4 calibration is retained. Timing includes initial prefill, generation, cache handling, scoring, final checking and orchestration. Model loading and warmup are recorded separately. All eight measured designs completed without checkpoint recovery.

**Table 12.** Full-workflow batch calibration, separate from the case-study outputs. The lowest complete time selects each backend’s batch (bold), using the same 3,072 bp M-pattern workload and calibration seed 2026091801. The native batch-4 output fails the predictive qualification threshold; quality is reported but does not select or exclude a batch.

| Backend | Batch | Complete time (min) | Checker AUROC |
| --- | --- | --- | --- |
| Native | 1 | 35.831 | 0.9653 |
| Native | 2 | 24.216 | 0.9692 |
| Native | 4 | 17.313 | 0.8841 |
| Native | 8 | **13.663** | 0.9501 |
| EvSpark | 1 | 12.188 | 0.9874 |
| EvSpark | 2 | 10.454 | 0.9631 |
| EvSpark | 4 | **8.509** | 0.9900 |

### Analysis and artifacts

The unit of analysis is a complete design (Table 13). The same two seeds are reused across targets; seed-group bootstrap intervals are exploratory and are not used to establish equivalence. We report all four paired time ratios. For the fixed-budget curve, each arm executes the frozen order M/seed 1, M/seed 2, S/seed 1, S/seed 2; only qualified, exact-unique top-ranked outputs completed within the common observed horizon are counted. Pairwise positionwise identity and pooled 4-mer entropy describe the four final sequences per backend, without treating sequence pairs as independent replicates. The artifact contains the reference sequences, target-track tables, model revision and weight SHA256 manifests, environment records, all final sequences, all search trajectories, batch calibration, and unrounded stage timings. CPU-only scripts audit the 5,640 candidate extensions and regenerate the figure from stored predictions, without loading models.

**Table 13.** Every complete case-study design. Native uses batch 8; EvSpark uses batch 4. M and S are the 768/768 bp and 384/1,152 bp open/closed targets. Seeds 1/2 are 2026091901/2026091902, reused across targets. All eight outputs meet both AUROC thresholds of 0.90. Search and checker columns report predictive AUROC, not experimental function.

| Pattern | Seed | Backend | Time (min) | Search | Checker |
| --- | --- | --- | --- | --- | --- |
| M | 1 | Native | 13.623 | 0.9703 | 0.9670 |
| M | 1 | EvSpark | 8.707 | 0.9796 | 0.9657 |
| M | 2 | Native | 13.662 | 0.9497 | 0.9457 |
| M | 2 | EvSpark | 8.038 | 0.9768 | 0.9774 |
| S | 1 | Native | 13.614 | 0.9988 | 0.9983 |
| S | 1 | EvSpark | 8.603 | 0.9806 | 0.9832 |
| S | 2 | Native | 13.620 | 0.9945 | 0.9884 |
| S | 2 | EvSpark | 9.587 | 0.9852 | 0.9821 |

## Supporting information

acceleration demo

## Footnotes

1 The post-prefill truncation behavior and a proposed guard are documented in https://github.com/Zymrael/vortex/issues/81 and https://github.com/Zymrael/vortex/pull/82.

2 Reproducer and environment details: https://github.com/Zymrael/vortex/issues/83.

## Notes

https://github.com/dhnihaoya/EvSpark

https://huggingface.co/dinghhhhhhhhhhhhhhh/EvSpark

### Competing Interest Statement

The authors have declared no competing interest.

### Summary of Updates

This revision clarifies methods and statistical reporting, replaces GPU-hour estimates with recorded wall-clock training times, and corrects figure labels and references. The author list and supplementary data are updated; the main conclusions remain unchanged.

## References

[1] Brixi, G., Durrant, M. G., Ku, J., Naghipourfar, M., Poli, M., Sun, G., et al. Genome modelling and design across all domains of life with Evo 2. Nature, 652(8112):1349–1361, 2026.

[2] Nguyen, E., Poli, M., Durrant, M. G., Kang, B., Katrekar, D., Li, D. B., et al. Sequence modeling and design from molecular to genome scale with Evo. Science, 386(6723):eado9336, 2024.

[3] Nguyen, E., Poli, M., Faizi, M., Thomas, A., Wornow, M., Birch-Sykes, C., et al. HyenaDNA: Long-Range Genomic Sequence Modeling at Single Nucleotide Resolution. NeurIPS, 36:43177–43201, 2023.

[4] Dalla-Torre, H., Gonzalez, L., Mendoza-Revilla, J., Lopez Carranza, N., Grzywaczewski, A. H., Oteri, F., et al. Nucleotide Transformer: building and evaluating robust foundation models for human genomics. Nature Methods, 22(2):287–297, 2025.

[5] Schiff, Y., Kao, C. H., Gokaslan, A., Dao, T., Gu, A., and Kuleshov, V. Caduceus: Bi-Directional Equivariant Long-Range DNA Sequence Modeling. ICML (PMLR 235), pp. 43632–43648, 2024.

[6] Shao, B. and Yan, J. A long-context language model for deciphering and generating bacteriophage genomes. Nature Communications, 15(1):9392, 2024.

[7] Provatas, K., Karatzikos, A., Koilakos, C., Patsakis, M., Tzanakakis, A., Nayak, A., et al. Accelerating inference in genomic and proteomic foundation models via speculative decoding. Bioinformatics, 42(8):btag579, 2026.

[8] Wu, Y., Dukler, Y., Trager, M., Achille, A., Xia, W., and Soatto, S. Snakes and Ladders: Accelerating SSM Inference with Speculative Decoding. NeurIPS ENLSP Workshop (PMLR 262), pp. 292–304, 2024.

[9] Wu, Y., Qin, Z., Wong, A., and Soatto, S. STree: Speculative Tree Decoding for Hybrid State Space Models. NeurIPS, 38:115588–115608, 2025.

[10] Wang, J., Paliotta, D., May, A., Rush, A. M., and Dao, T. The Mamba in the Llama: Distilling and Accelerating Hybrid Models. NeurIPS, 37:62432–62457, 2024.

[11] Cheng, X., Yu, X., Shao, C., Li, J., Xiong, Y., Qian, Y., et al. DSpark: Confidence-Scheduled Speculative Decoding with Semi-Autoregressive Generation. 2607.05147, 2026.

[12] Poli, M., Wang, J., Massaroli, S., Quesnelle, J., Carlow, R., Nguyen, E., et al. StripedHyena: Moving Beyond Transformers with Hybrid Signal Processing Models. GitHub software repository, 2023. https://github.com/togethercomputer/stripedhyena.

[13] Poli, M., Massaroli, S., Nguyen, E., Fu, D. Y., Dao, T., Baccus, S., et al. Hyena Hierarchy: Towards Larger Convolutional Language Models. ICML (PMLR 202), pp. 28043–28078, 2023.

[14] Leviathan, Y., Kalman, M., and Matias, Y. Fast Inference from Transformers via Speculative Decoding. ICML (PMLR 202), pp. 19274–19286, 2023.

[15] Chen, C., Borgeaud, S., Irving, G., Lespiau, J.-B., Sifre, L., and Jumper, J. Accelerating Large Language Model Decoding with Speculative Sampling. 2302.01318, 2023.

[16] Stern, M., Shazeer, N., and Uszkoreit, J. Blockwise Parallel Decoding for Deep Autoregressive Models. NeurIPS, 31, 2018.

[17] Zhang, D., Zhang, W., Zhao, Y., Zhang, J., He, B., Qin, C., et al. DNAGPT: A Generalized Pre-trained Tool for Versatile DNA Sequence Analysis Tasks. 2307.05628, 2023.

[18] Nijkamp, E., Ruffolo, J. A., Weinstein, E. N., Naik, N., and Madani, A. ProGen2: Exploring the boundaries of protein language models. Cell Systems, 14(11):968–978.e3, 2023.

[19] Ferruz, N., Schmidt, S., and Höcker, B. ProtGPT2 is a deep unsupervised language model for protein design. Nature Communications, 13(1):4348, 2022.

[20] Li, Y., Wei, F., Zhang, C., and Zhang, H. EAGLE: Speculative Sampling Requires Rethinking Feature Uncertainty. ICML (PMLR 235), pp. 28935–28948, 2024.

[21] Cai, T., Li, Y., Geng, Z., Peng, H., Lee, J. D., Chen, D., et al. Medusa: Simple LLM Inference Acceleration Framework with Multiple Decoding Heads. ICML (PMLR 235), pp. 5209–5235, 2024.

[22] Ankner, Z., Parthasarathy, R., Nrusimha, A., Rinard, C., Ragan-Kelley, J., and Brandon, W. Hydra: Sequentially-Dependent Draft Heads for Medusa Decoding. COLM, 2024.

[23] Li, Y., Wei, F., Zhang, C., and Zhang, H. EAGLE-2: Faster Inference of Language Models with Dynamic Draft Trees. EMNLP, pp. 7421–7432, 2024.

[24] Zhou, Y., Lyu, K., Rawat, A. S., Menon, A. K., Rostamizadeh, A., Kumar, S., et al. DistillSpec: Improving Speculative Decoding via Knowledge Distillation. ICLR, pp. 32011–32050, 2024.

[25] Miao, X., Oliaro, G., Zhang, Z., Cheng, X., Wang, Z., Zhang, Z., et al. SpecInfer: Accelerating Large Language Model Serving with Tree-based Speculative Inference and Verification. ASPLOS, volume 3, pp. 932–949, 2024.

[26] Fu, Y., Bailis, P., Stoica, I., and Zhang, H. Break the Sequential Dependency of LLM Inference Using Lookahead Decoding. ICML (PMLR 235), pp. 14060–14079, 2024.

[27] Saxena, A. Prompt Lookup Decoding. GitHub software repository, 2023. https://github.com/apoorvumang/prompt-lookup-decoding.

[28] Gu, A. and Dao, T. Mamba: Linear-Time Sequence Modeling with Selective State Spaces. COLM, 2024.

[29] Lieber, O., Lenz, B., Bata, H., Cohen, G., Osin, J., Dalmedigos, I., et al. Jamba: A Hybrid Transformer-Mamba Language Model. 2403.19887, 2024.

[30] Parks, D. H., Chuvochina, M., Rinke, C., Mussig, A. J., Chaumeil, P.-A., and Hugenholtz, P. GTDB: an ongoing census of bacterial and archaeal diversity through a phylogenetically consistent, rank normalized and complete genome-based taxonomy. Nucleic Acids Research, 50(D1):D785–D794, 2022.

[31] Camargo, A. P., Nayfach, S., Chen, I.-M. A., Palaniappan, K., Ratner, A., Chu, K., et al. IMG/VR v4: an expanded database of uncultivated virus genomes within a framework of extensive functional, taxonomic, and ecological metadata. Nucleic Acids Research, 51(D1):D733–D743, 2023.

[32] Avsec, Ž., Agarwal, V., Visentin, D., Ledsam, J. R., Grabska-Barwinska, A., Taylor, K. R., et al. Effective gene expression prediction from sequence by integrating long-range interactions. Nature Methods, 18(10):1196–1203, 2021.

[33] Linder, J., Srivastava, D., Yuan, H., Agarwal, V., and Kelley, D. R. Predicting RNA-seq coverage from DNA sequence as a unifying model of gene regulation. Nature Genetics, 57(4):949–961, 2025.

[34] Hingerl, J. C., Karollus, A., and Gagneur, J. Flashzoi: an enhanced Borzoi for accelerated genomic analysis. Bioinformatics, 41(9):btaf467, 2025.

