## Supplementary figures and images for "EvSpark: Lossless Speculative Decoding for Hybrid DNA Foundation Models"

### acceleration demo

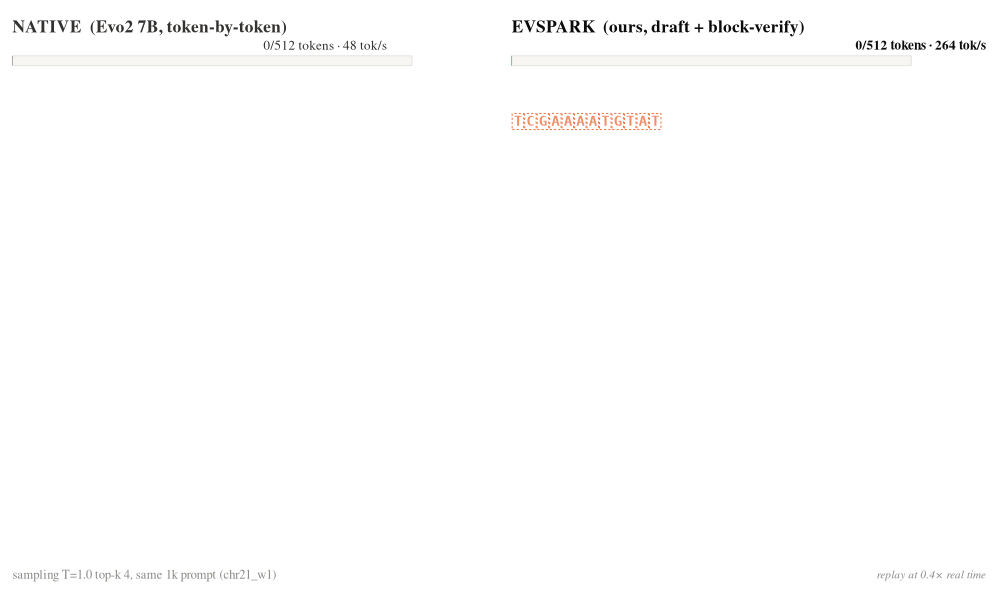
